# A large-scale optogenetic neurophysiology platform for improving accessibility in non-human primate behavioral experiments

**DOI:** 10.1101/2024.06.25.600719

**Authors:** Devon J. Griggs, Noah Stanis, Julien Bloch, Jasmine Zhou, Karam Khateeb, Shawn Fisher, Larry Shupe, Eberhard E. Fetz, Hesamoddin Jahanian, Azadeh Yazdan-Shahmorad

## Abstract

Optogenetics has been a powerful scientific tool for two decades, yet its integration with non-human primate (NHP) electrophysiology has been limited due to several technical challenges. These include a lack of electrode arrays capable of supporting large-scale and long-term optical access, inaccessible viral vector delivery methods for transfection of large regions of cortex, a paucity of hardware designed for large-scale patterned cortical illumination, and limited designs for multi-modal experimentation. To address these gaps, we introduce a highly accessible platform integrating optogenetics and electrophysiology for behavioral and neural modulation with neurophysiological recording in NHPs. We employed this platform in two rhesus macaques and showcased its capability of optogenetically disrupting reaches, while simultaneously monitoring ongoing electrocorticography activity underlying the stimulation-induced behavioral changes. The platform exhibits long-term stability and functionality, thereby facilitating large-scale electrophysiology, optical imaging, and optogenetics over months, which is crucial for translationally relevant multi-modal studies of neurological and neuropsychiatric disorders.

**Graphical Abstract:** 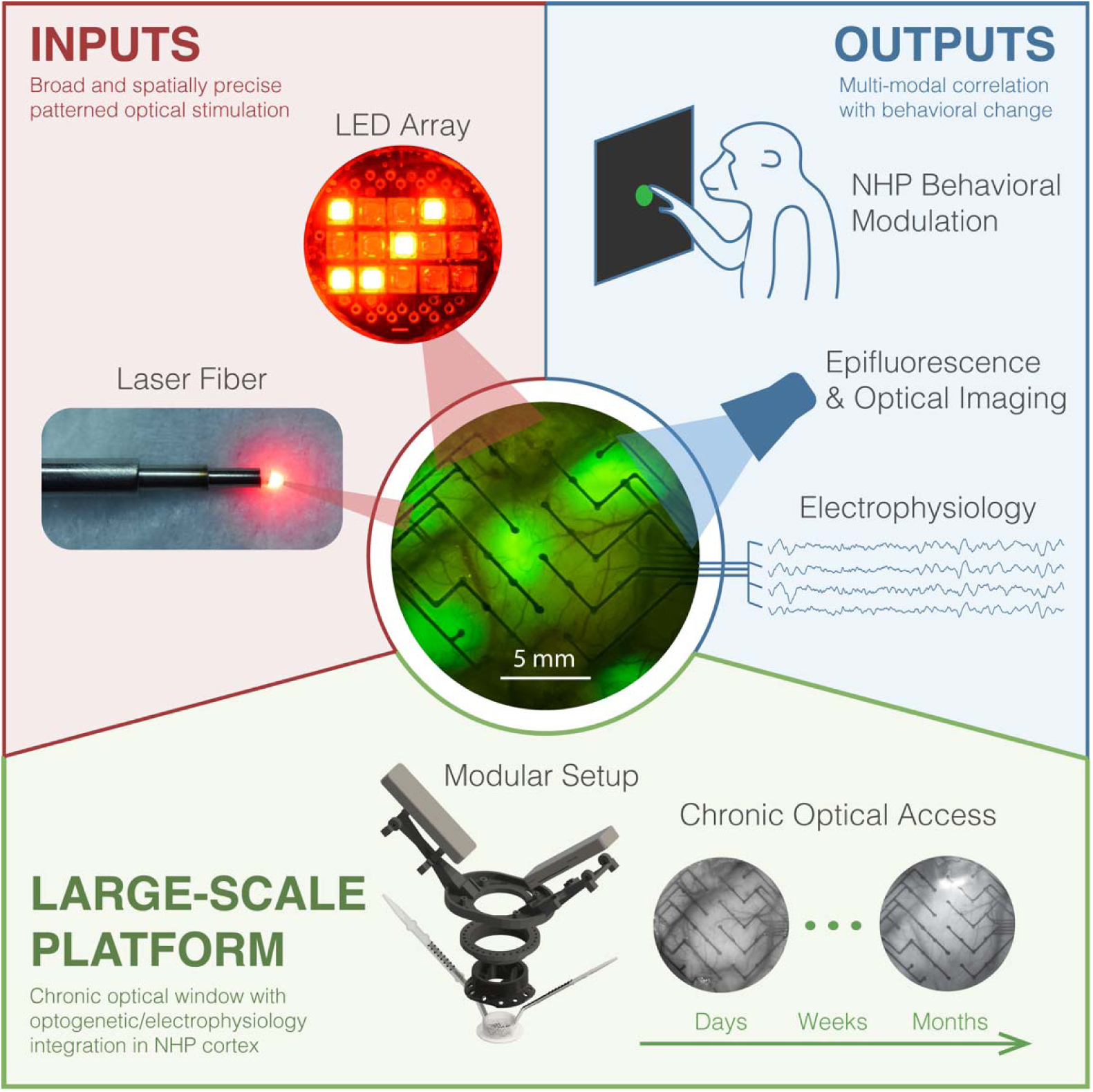

## Introduction

Optogenetics is unique among neuromodulation technologies because it allows for cell-type specific manipulation of neural networks with high spatial and temporal precision. This enables the precise manipulations needed for research in fundamental neuroscience ^1–5^, such as investigating relationships between neural activity and behavior ^4–8^, monitoring changes in neural circuitry over time ^7,9^, understanding the neural changes associated with brain dysfunction ^10,11^ and developing therapies for neurological disorders ^1,3,5,12,13^.

The evolutionary proximity between the neurophysiology of non-human primates (NHPs) and humans ^14,15^ spurs us to apply optogenetics to NHPs to better understand the human brain and strengthen translational medical research, especially in the context of neurological and neuropsychiatric disorders. However, studying neurological disease models in NHPs is challenging because NHP neuropathology and anomalous neural activity are often spatiotemporally complex, spanning multiple brain areas and months or years of development, much like in humans. These challenges, along with the scientific opportunities afforded by optogenetics, call for the development of large-scale, chronically-stable optogenetic platforms for NHPs. Although the development of optogenetic techniques in NHPs emerged soon after the debut of optogenetics ^16,17^, its wide-spread adoption has remained a challenge. We dissect key contributing factors below that are addressed in our platform.

While optogenetics alone is a potent technique for functional neuropathological studies, the integration of optogenetics with an electrical platform, such as electrocorticography (ECoG) recordings, can enable precise optical control over specific cell types with concurrent measurement of neural activity at high spatiotemporal resolution. Merging of optogenetic and electrical recording platforms has been demonstrated in rodent work ^18,19^, however, similar attempts in NHP work have come with limitations. Systems where skin is closed over the implant require surgeries for device replacements or modifications and prevent ongoing assessment of both optical occlusions between the light source and the brain, as well as optogenetic expression ^20^, which is typically confirmed with optical techniques. Such limitations make these systems experimentally inflexible.

In contrast, modular chambers enabling chronic access to the cortical surface have been developed. In particular, *in vivo* optical imaging of NHP brains through cranial windows has been a small but vibrant field with decades of development with techniques such as calcium imaging^21^, voltage sensitive dyes^22,23^, intrinsic signal optical imaging^23,24^, and epifluoresence imaging^9,25^, among others. Optical windows to the NHP brain are often covered with a transparent, silicone-based artificial dura to protect the brain while providing optical access to the cortex ^22–25^. The artificial dura has been combined with optical stimulation methods such as infrared neural stimulation ^26^ and optogenetics ^24,27^. In a previous effort, we have taken a similar approach and performed simultaneous electrical recording and optogenetic stimulation using a µECoG array in an optical window. However, long-term optical access stability using this approach remained an unmet challenge. One approach relied on daily explantation of an artificial dura and implantation of a µECoG array to perform experiments, but the frequent manipulation of the cortical surface stimulated tissue growth over the cortex and obscured optical access ^25,28^. Another approach relied on chronic co-implantation of both an artificial dura and a µECoG array between the artificial dura and the brain, but tissue growth between the artificial dura and the µECoG array obscured optical access ^28,29^. To take full advantage of optogenetics in NHPs, there is a need for innovative platforms with chronic stability in both electrophysiology and optical access to empower widespread adoption.

One obstacle in NHP optogenetics, not present in rodent models, is the size of NHP brains, which renders widespread viral delivery of optogenetic vectors impractical via traditional, diffusion-based methods. In contrast, convection-enhanced delivery (CED) is a pressure-based approach of infusion allowing for large-scale and even distribution of viral expression in comparison to diffusion-based approaches ^25,30^. Previously published CED protocols relied on live magnetic resonance imaging (MRI) to monitor infusions ^31,32^, making these methods technically challenging and only accessible to the institutions capable of MRI-guided viral infusions in NHP brain.

Given the challenges of attaining large-scale optogenetic coverage and maintaining optical access, large-scale optical stimulation techniques have been largely undeveloped – optogenetic stimulation in NHPs is generally performed by fiber optics or optrodes coupled to lasers ^33^, which are not optimized for large-area optogenetic coverage. Light emitting diode (LED) arrays, both with ^20^ and without ^34,35^ simultaneous electrophysiological recording, have also been used, but with limited cortical coverage or access. Optogenetic transduction and stimulation methods, that are both experimentally flexible and widely accessible, are necessary to capitalize on large-scale optogenetics in NHPs.

In this paper, we address the above challenges by designing accessible tools and integrating them into a single platform. We demonstrate large-scale, chronically-stable optogenetics and ECoG in an imaging window on NHP cortex. We present an optogenetic platform comprised of five complimentary state-of-the-art technological advancements, enhancing both experimental flexibility and accessibility to researchers with diverse technical backgrounds. Our advancements are: (1) a platform designed to be modular and flexible for varied, multi-modal use-cases; (2) a chronically implanted integration of an ECoG array and an artificial dura, which we refer to as a multi-modal artificial dura (MMAD); (3) optical hardware scalable from single lasers up to large arrays of LEDs for complex patterned stimulation across multiple brain areas; (4) improved ECoG-compatible optical access with the MMAD, facilitating long-term multi-modal experimentation; and (5) CED without live-MRI guidance, which is an accessible viral vector delivery method for achieving large-scale expression. These technologies are synergistic yet maintain functional modularity, i.e., no one technology requires all the others to function, thus keeping our platform relevant for experiments requiring fewer than our full suite of technologies, while maintaining forward compatibility. Our work is poised to bridge the gap between rodents and humans by facilitating powerful, multi-modal studies in NHPs.

We demonstrated the efficacy of our advancements by three complimentary measures: optical imaging, electrophysiology, and behavior. We present months of continuous optical clarity with the MMAD and years of chamber stability, which opens opportunities for translational studies of models of neurological diseases and disorders such as stroke recovery ^36^. The optical clarity empowered verification of large-scale epifluorescence, the product of our viral delivery process, and our ECoG recordings confirmed the flexibility of our scalable optical stimulation setup for optogenetic experimentation. We demonstrated our platform’s relevance for behavioral studies by optogenetically disrupting a center-out reach task by modulating the posterior parietal cortex (PPC). This showcases the applicability of our platform for complex behavioral experiments. This work provides a milestone proof-of-concept for NHP behavior experiments with inhibitory opsins which, up to this point, have not been successfully applied to reaching behavior.

The ability to link millisecond-resolution neural activity with behavioral effects and optogenetic neuronal manipulation is an invaluable addition to the NHP researcher’s toolbox. Therefore, we took measures to ensure the accessibility of our methods for a diverse range of specialties: design files are provided for parts and equipment that are not commercially available, our in-house fabrication processes require only generic equipment and modest skill, and our neural processing code is publicly available. Collectively, our work enables large-scale and long-term interrogation of cortical areas in awake behaving NHPs ^37,38^.

## Results

### Hardware

We designed and implemented our platform for two healthy adult male rhesus macaques (*Macaca mulatta*, Monkeys H and L). We used magnetic resonance imaging (MRI)-based methods to plan a craniotomy with a diameter (Ø) of 25 mm over the left PPC and to design a custom chamber with a curvature precisely fitting the surface of the skull around the craniotomy for each monkey (Fig. 1a, c). We have previously published these methods as step-by-step protocols for accessibility ^39,40^. The custom chamber was designed to provide a sturdy base for experimental equipment and protect the brain when the monkey is freely moving between experiments. The chamber is further described in STAR Methods. We implanted the chamber along with our multi-modal artificial dura (MMAD; Fig. 1b, d; ^41^) in both animals. The chambers continue to be stable and used for studies after >4.0 and >3.3 years following implantation for Monkeys H and L, respectively, and we have not observed behavioral or health deficits.

**Figure 1:**
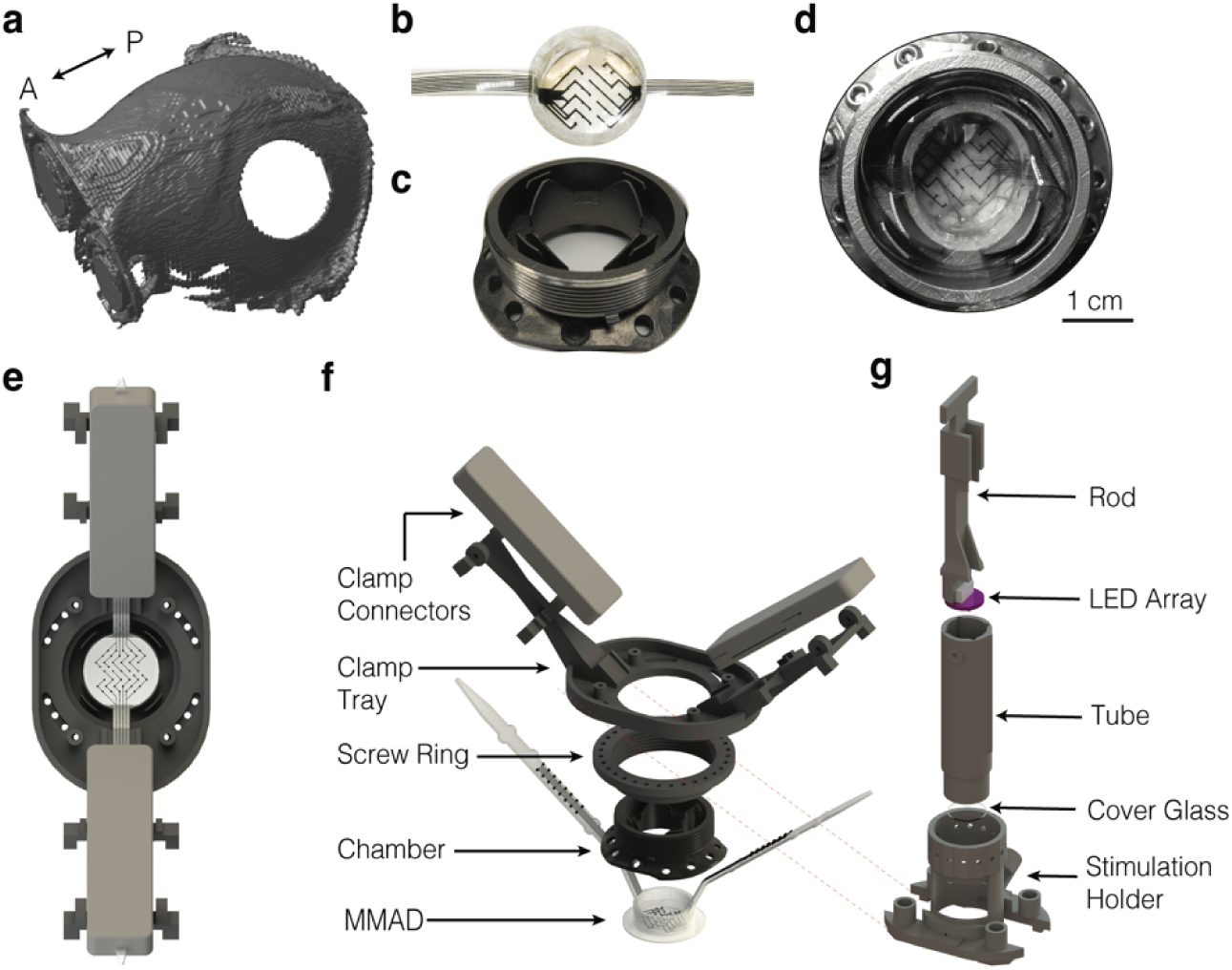
Hardware and skull model. **(a)** Digital representation of the skull with craniotomy. **(b)** Molded MMAD. **(c)** Chamber custom-fit to the skull. **(d)** Implanted chamber and MMAD packaged inside while cables are housed in the concentric grooves. **(e)** Model of electrophysiological recording hardware, top view and **(f)** trimetric view. From bottom up: MMAD, chamber, ring, clamp connector tray, clamp connectors. **(g)** LED-based optical stimulation hardware. From bottom up: Stimulation holder, cover glass, tube, LED array, rod. Panels (a, f, g) are adapted from^39^ with permission from IEEE EMBC.

We designed the MMAD to serve both as an optical window to the brain and as an electrocorticography (ECoG) array. The MMAD is composed of a flexible and corrosion resistant electrode array molded in silicone into a “top-hat” shape common for traditional artificial duras ^40,42^. The opaque, conductive traces of the MMAD are layered on top of each other to maximize the optical access to the brain through the transparent polymer of the MMAD ^41^. Like traditional artificial duras, the MMAD protects the brain and keeps the brain moist during experiments. Unlike traditional artificial duras, cables extend from the “top-hat” of the MMAD to hardware for electrophysiological recordings, as described below. We packaged the MMAD cables inside the chamber between experiments (Fig. 1d). The MMAD, as well as the commercially available electrode array (Ripple Neuro, Salt Lake City, UT) from which the MMAD was fabricated, are further described in STAR Methods.

The MMAD cables require adapters, called clamp connectors, to be used with external electrophysiology equipment. We designed a stack-up of parts to attach to the chamber and support the clamp connectors (Fig. 1e, f). All parts are designed to be modular for flexible experimentation and design iterations as needed. Further descriptions of parts can be found in STAR Methods, and design files are publicly available (see Data and Code Availability).

We designed and tested our platform for both laser- and LED-based optical stimulation. The lasers and optical fibers were commercially sourced and can support a variety of light wavelengths (see STAR Methods for further laser system details). We designed an LED array and its driving circuitry in-house (Fig. 2, Supp. Fig. 1) to be compatible with high temporal resolution (sub-millisecond) requirements. While pulse-width modulation is a popular technique for driving LEDs, we designed our driving circuitry to be controlled by analog signals. Specifically, we drove our LEDs with continuous voltage signals to avoid high-frequency photo-induced artifacts in our electrophysiology recordings. We selected LEDs for optimal tissue penetration and powered our circuitry with two rechargeable 9V batteries to mitigate line noise (Supp. Fig. 1). We designed and fabricated two versions of an LED array for patterned stimulation, a 4 × 4 array with individually drivable LED columns ^43^ and a 3 × 5 array with independently drivable LEDs (Fig. 2a). An example stimulation pattern demonstrating spatial, temporal, and optical power complexity is provided (Supp. Vid. 1). Both boards used identical schematics for the driving circuitry except for the number of LEDs driven by each driving circuit (Supp. Fig. 1). We fabricated our LED arrays in two wavelengths (634 nm and 485 nm). The LED array was protected from potential moisture from the brain by a cover glass between the LED array and the MMAD (Fig. 1e-g, 2b). This also provided a small air gap between the LEDs and the cover glass to mitigate conductive heating of the brain (Fig. 2b). Our benchtop temperature measurement data indicate our stimulation design limits the cortical temperature change to less than 1 ℃ during our stimulation protocols (Supp. Fig. 2), which is below the safe limit for brain ^44–47^ and has minimal impact on neuronal activity ^48,49^. We did not observe any tissue damage (e.g., brain discoloration) or behavioral deficits over the years in our animals. We commercially sourced our custom printed circuit boards (PCBs) and assembled the LED array and driver PCBs in-house with common soldering methods. (For more details, see STAR Methods.)

**Figure 2:**
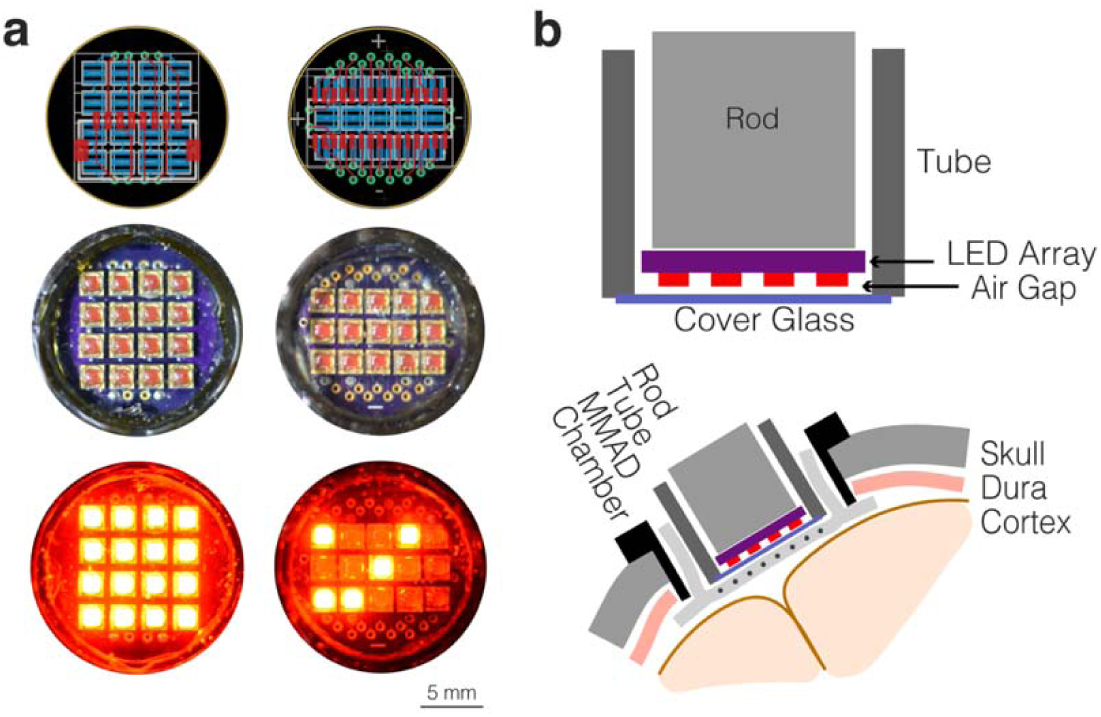
LED array. **(a)** LED column-drivable 4 × 4 LED array (left column) and individually drivable 3 × 5 LED array (right column). PCB schematic (top), photograph of LED array in tube and through the cover glass with LEDs off (middle) and on in an example illumination configurations (bottom). **(b)** Schematic of LED array and related hardware for optical stimulation through MMAD, including an air gap to mitigate tissue heating. Upper- and middle-left panels of (a) are adapted from^39^ with permission from IEEE EMBC.

### Optical stability

To demonstrate the stability of optical access through the MMAD we used both optical epifluorescence imaging and stimulation. With our MMAD, we achieved 10 to 14 consecutive weeks of optical access in our monkeys before tissue growth began to obstruct light penetration (Supp. Fig. 3). After tissue had matured, we resected the tissue and reimplanted the MMAD for additional experiments.

As aforementioned, both implanted monkeys have been stable for ∼4 and ∼3 years following surgery, and the chambers have supported optical experiments, displaying the platform’s ability to facilitate long-term NHP studies. Over this period, we have repeated the dura resection and reimplantation of the MMAD for both monkeys, each allowing us windows of >10 weeks of optical access for experimentation. In some instances, we observed accelerated tissue growth when we explanted the MMAD to test other recording and stimulation systems. For example, following a tissue resection in Monkey H approximately two years after first implantation, the MMAD was replaced and clear optical access was maintained for 7 weeks. At week 7, while optical access was still clear, the MMAD was removed to test other stimulation and recording devices that impacted tissue growth. Approximately one week later, tissue began to grow over the cortex. For another example, one and a half years after first implantation in Monkey L, a similar procedure provided clear optical access for at least 5.5 weeks during which we ran experiments. Similarly, the MMAD was removed for another experiment that led to tissue growth. These results collectively show the stability of optical access with our design for several months and the modularity of our platform to test other recording and stimulation devices. In addition, our results caution that explantation and implantation of the MMAD must be minimized, because it is known that mechanical manipulation on the surface of the brain causes irritation of the cortical surface and can lead to accelerated tissue growth that obscures optical access.

### CED and epifluorescence

During the chamber implantation surgery, we used CED (similar to ^9,25,50,51^) to infuse up to 50 µL of a pan-neuronal inhibitory optogenetic viral vector (AAV8-hSyn-Jaws-GFP, 5.4 x 10^12^ genome copies per milliliter (gc/mL)) ^52,53^ at several locations across the craniotomy (Fig. 3). We delivered viral vector at rates up to 5 µL/min using our custom-built stepped-tip cannula ^25^. Further descriptions of parts, equipment, and protocols are provided in STAR Methods.

**Figure 3:**
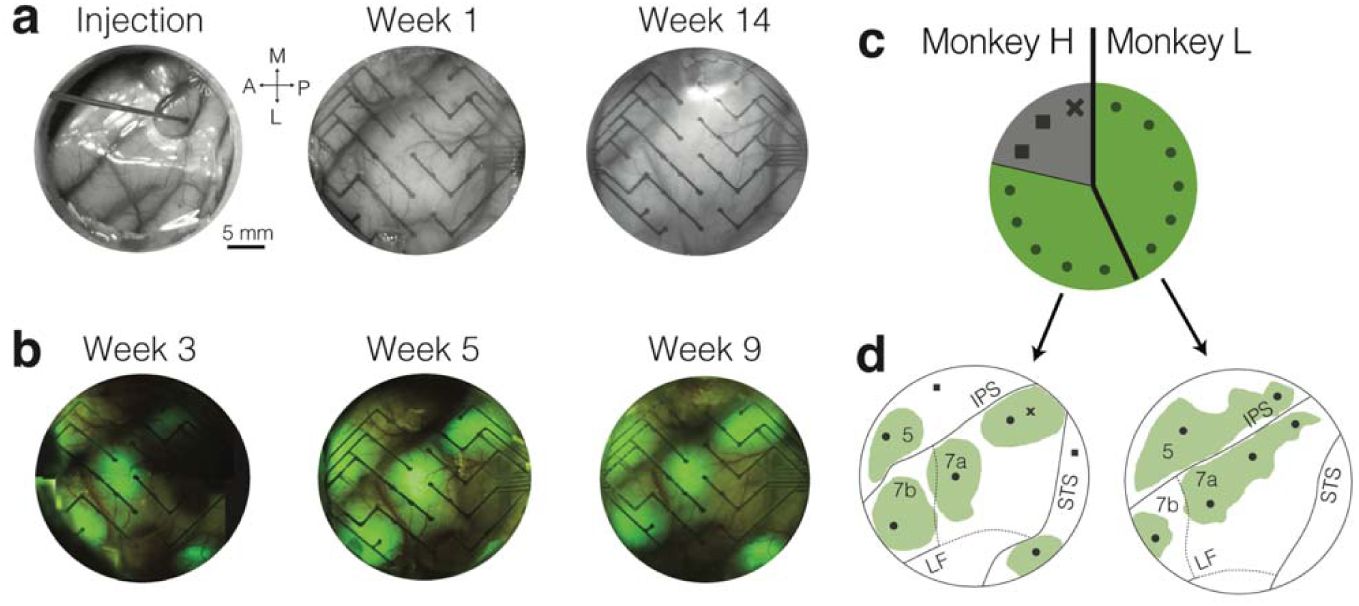
Optical window and epifluorescence. **(a)** Optical access and **(b)** epifluorescence for Monkey H over time. CED was performed through a hole in an artificial dura during surgery to virally transfect tissue (a, left). **(c)** Pie-graph summary of CED infusions across both monkeys, which highlights the success of our methodological improvement in our second surgery (Monkey L). Successful infusions are grey dots and unsuccessful infusions are grey squares and one infusions canceled before completion is represented by a grey ×. **(d)** Expression maps based on epifluorescence (green) for Monkeys H (left, first monkey to undergo surgery) and L (right, second monkey to undergo surgery). Approximate locations of successful (green dots) and unsuccessful (grey squares) CED infusions. One infusion was canceled before completion (grey ×) and a second infusion attempt was made nearby, which was successful. Portions of this figure (a, center; a, right; b, center; and d, left) are adapted from39 with permission from IEEE EMBC.

We infused the optogenetic viral vector in the surgery suite immediately prior to chamber implantation. This allowed us to visually inspect any potential reflux on the surface of the brain from the cannula without live-MRI, which we had used in previous protocols. For this new protocol, we had eleven successful and two unsuccessful infusions as determined by epifluorecent expression near the locations of cannula insertion. We canceled one infusion before its completion because the brain did not seal against the cannula and we observed fluid buildup, likely reflux (Fig. 3c, d). We reattempted that infusion nearby and successfully generated expression (Fig. 3d). In the first surgery (Monkey H), we had difficulty determining the depth of cannula penetration into the brain with stereotactic instruments alone because of the brain’s tendency to dimple during cannula insertion. To address this in our second surgery (Monkey L), we marked the cannula prior to surgery at a set distance distal from the cannula’s tip to facilitate measurement of penetration depth during surgery. This technique correlated with an improvement in success rate between our first and second surgeries (62% and 100%, respectively; Fig. 3c, d).

Our long-term and large-scale optical access provided by the MMAD empowered us to monitor optogenetic expression over time for both monkeys. Monkey H underwent initial imaging for epifluorescence two weeks after viral infusion. Expression was evident during the first imaging session and increased in coverage and intensity until the fifth week after infusion, when expression plateaued at 92 mm^2^ coverage (Fig. 3b, d). Similarly, we confirmed 76 mm^2^ expression coverage in Monkey L (Fig. 3d). Although we did not see a change in epifluorescent expression during the initial months following the plateau in viral expression area, approximately two years after infusion we observed a strong reduction of expression in both monkeys.

### Photo-induced artifact removal

Our optical stimulation system generated artifacts in the ECoG data. To remove artifacts from ECoG data, we generated a dataset of artifacts, in the absence of any neural signals, by optically stimulating an MMAD submerged in a saline bath. We modeled the photo-induced artifact using these saline data, fit the model to the *in vivo* ECoG data, and removed the estimated photo-induced artifact (see Methods for details). The results demonstrate that the method effectively removes photo-induced artifacts (Fig. 4a). As a control, we tested the artifact removal method on data without stimulation and found that the method had negligible effect, as intended (Fig 4b, c).

**Figure 4:**
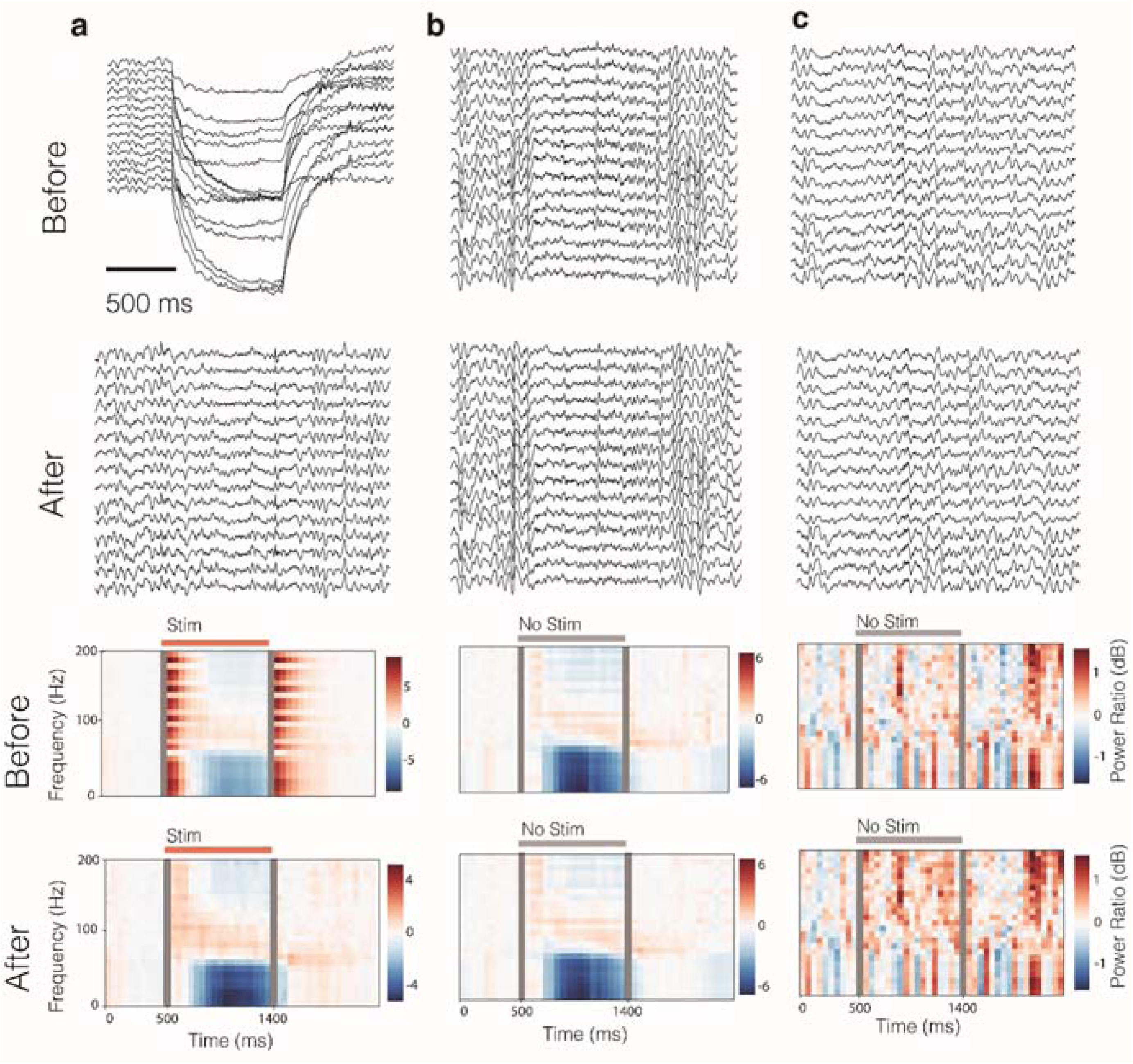
Removal of photo-induced artifact. Traces are from before (first row) and after (second row) artifact removal. Likewise, spectrograms are from before (third row) and after (fourth row) artifact removal. Example traces and spectrograms of **(a)** optical stimulation of the cortex when the animal was performing a reach, **(b)** no optical stimulation when the animal was performing a reach, and **(c)** no optical stimulation when the animal was at rest. Note that the time-scale differs between traces and spectrogram, and note that the scale of color bars differs between spectrograms, particularly for the at-rest condition. Grey bars on spectrograms indicate bins during the first 50 ms of stimulation onset and offset – data during these times are excluded due to residual artifact after the artifact removal process.

### Stimulation at rest

To demonstrate our platform’s ability to integrate optogenetics and ECoG, we applied optical stimulation protocols (Fig. 5a, b) and recorded evoked neural activity. We removed photo-induced artifact as described above and in STAR Methods. Statistical tests were run by comparing the neural activity to the baseline recording before the onset of stimulation (Bonferroni correction applied to p < 0.05).

**Figure 5:**
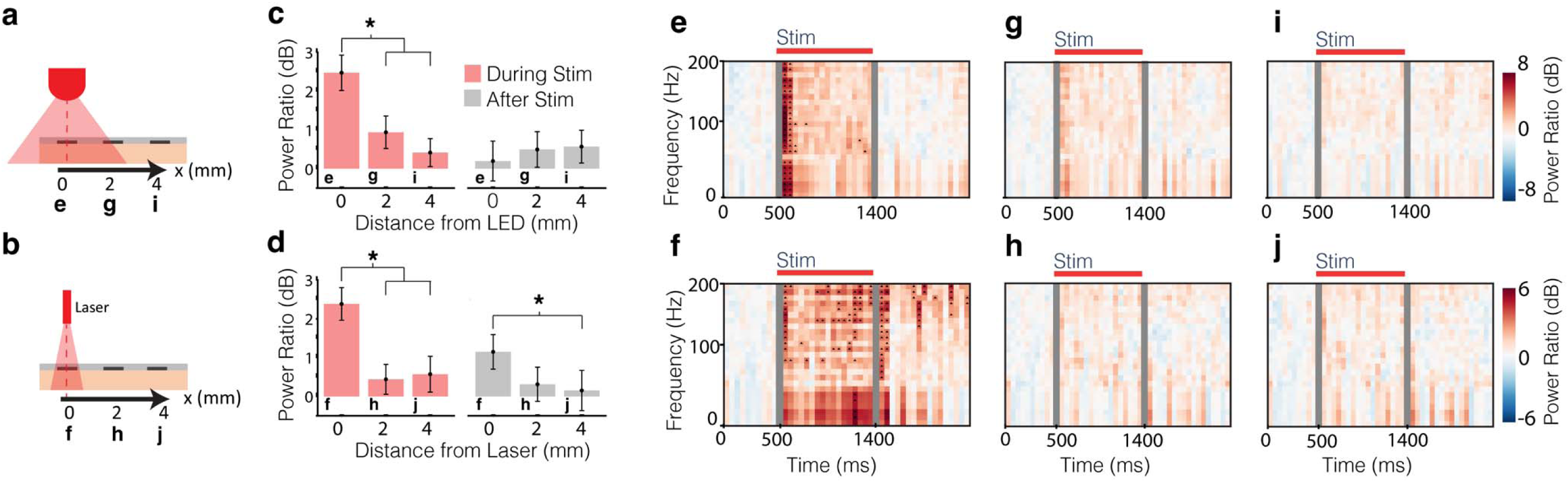
The effect of stimulation on neural activity during rest. **(a, b)** Schematics of optical stimulation protocols. (a): single red LED (634 nm) with recording electrodes at varying distances from the light source (e, g, i). (b): single red laser (638 nm) with recording electrodes at varying distances from the light source (f, h, j). **(c, d)** The power ratios of responses are calculated by taking the ECoG power during stimulation (red bars), or after stimulation (grey bars), and dividing by the ECoG power from before stimulation (1-200 Hz), then converting to decibels. Whiskers are 95% confidence intervals of the medians generated with a bootstrapping method, and asterisks indicate p < 0.05 using the Mann-Whitney U test. **(e-j)** Spectrograms are power ratios where each time-frequency band was normalized to that band’s power in the 500 ms preceding stimulation. Spectrograms were averaged over 1 electrode and 30 trials for each location. The callouts at the base of the bars indicate the subpanel represented in that specific bar of the bar chart. On spectrograms, arrowheads indicate significant (Bonferroni correction applied to p < 0.05) differences between the time-frequency bin power and baseline power in that respective frequency band. Each stimulation offset was followed by 5 s before the following stimulation onset. Grey bars indicate bins during the first 50 ms of stimulation onset and offset – data during these times are excluded due to residual artifact after artifact removal. All data from Monkey L.

We stimulated an opsin-expressing area of Monkey L during rest with one of the red LEDs of the 3 × 5 LED array (634 nm, 75 mW, 900 ms; Fig. 5a). The stimulation produced a significant increase in ECoG power across most frequencies and weaker effects persisted after the end of stimulation (Fig. 5a, c, e) probably due to spatiotemporal complexities of neural networks. The effects of stimulation were similar but weaker at nearby electrodes (Fig. 5a, c, g, i). In addition to testing our LED array, we also tested traditional laser stimulation. We stimulated neural tissue of Monkey L with fiberoptic cable connected to a laser (638 nm, 45 mW, 900 ms; Fig. 5b). Again, the stimulation produced a significant increase in ECoG power across high frequencies and weaker effects persisted after the end of stimulation (Fig. 5b, d, f). Effects were primarily local (Fig. 5b, d, f, g, j), which aligned with our expectation that laser stimulation would have a more focal effect than LED stimulation. A similar experiment with Monkey H produced similar results (Supp. Fig. 4).

However, we expected that inhibitory optogenetics would cause ECoG power to decrease in general, not increase. To validate the light-evoked responses observed, we have performed a series of control experiments described below.

We investigated the spatial nature of the neural responses with respect to opsin expressing and opsin deficient areas (Fig. 6). To test the spatial nature of the neural responses, we stimulated all 16 red LEDs of the 4 × 4 array (634 nm, ∼12 mW/mm^2^) during rest and separated recording channels based on the expression in their vicinity. A control animal, Monkey C, which had never been genetically altered, did not show statistically significant changes in neural activity (*p* = 0.33; Fig. 6d). For both optogenetic monkeys, ECoG power evoked by red LEDs was statistically stronger in the opsin expressing areas than the opsin deficient areas (Monkey H: *p* = 0.012; Monkey L: *p* = 2.9e-9; Fig. 6a, c). We repeated the experiment with sub-optimal stimulation techniques in Monkey L, being red LED stimulation with tissue growth over the brain and blue (485 nm) LED stimulation of a clear window, and found there was no significant difference between the opsin expressing and opsin deficient areas (tissue: *p* = 0.44; blue light: *p* = 0.80; Fig. 6c). Repeating the red light stimulation experiment with the full LED array and clear optical access in Monkey H ∼2 years after initial infusion, when most of the epifluorescence had dissipated, produced no statistically significant changes from natural fluctuations in a baseline activity without optical stimulation (*p* = 0.89; Fig. 6b). Comparing light-evoked responses from opsin expressing and opsin deficient regions across days yielded statistically significant differences in Monkey L (Fig. 6e). Note that after 540 days we saw decreased expression as measured by both epifluorescence and insignificant light-evoked responses (*p* = 0.54; Fig. 6e). We then reinjected 838 days after initial injection (Fig. 6e) and generated approximately the same stimulation-evoked activity levels as the first injection in both opsin expressing and opsin deficient areas (Fig. 6e). Collectively, the differences between light-evoked responses of opsin expression and opsin deficient regions, the light evoked responses after reinjection, and the lack of response in control experiments (a naïve monkey, tissue growth, blue light) support our finding that optical stimulation of neural tissue expressing Jaws increased ECoG power, despite Jaws being an inhibitory opsin. This increase could be due to disinhibition of neurons in lower cortical layers following inhibition of upper cortical layers – we reason the disinhibition is reflected in the ECoG signal through apical dendrites from the lower layers.

**Figure 6:**
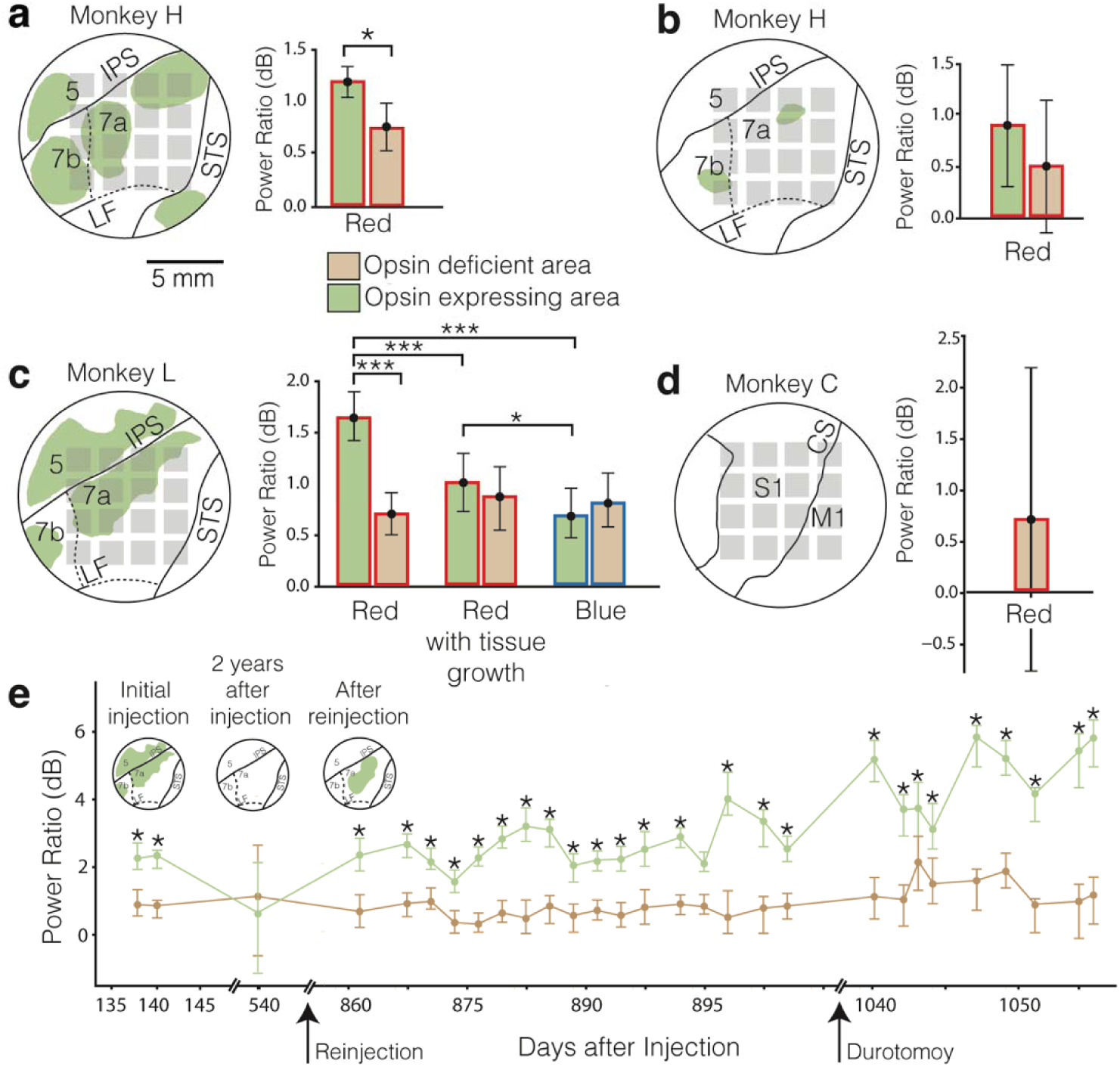
Neural responses with respect to opsin expressing and opsin deficient areas. **(a-d)** Epifluorescent maps and approximate LED locations for (a), Monkey H, 5 weeks after viral infusion, (b) Monkey H ∼2 years after viral infusion, (c) Monkey L, 4 weeks after viral infusion, and (d) Monkey C, a control animal with no viral infusion. Median evoked responses for each experiment are shown for light stimulation using the full 4×4 LED array. For Monkey L, evoked responses are also shown under control stimulation conditions of red light with tissue growth and blue light with clear optical access. The power ratios of responses are calculated by taking the ECoG power during stimulation and dividing by the ECoG power from before stimulation. The stimulation protocol was a pulse train of 900 ms stimulation followed by 5 s rest. Whiskers are 95% confidence intervals of the median generated with a bootstrapping method. Asterisks indicate significant differences between conditions (p < 0.05, Mann-Whitney U-test). **(e)** Median evoked responses in opsin deficient and expressing areas for Monkey L over time after initial infusion (left) and after reinjection 838 days later, including results following a durotomy of tissue growth 190 days after reinfusion (right). Opsin expression maps are shown for the initial injection, 2 years after injection, and after reinjection (inset). Trial counts for (a-d) can be found in Supp. Table 3.

Given the increase of ECoG power when inhibition was applied in vivo, we examined the potential underlying mechanism by performing a simulation experiment, investigating the effect of stimulation across different cortical depths. We used VERTEX ^54^, a simulation tool based on detailed cortical neuronal data ^55^ used for modeling and simulating large-scale networks of neurons, which we recently updated to support optogenetic stimulations ^56^. We found that pan-neuronal stimulation of Jaws-expressing neurons led to decreased spiking in upper layers (L2/3 and L4) and increased spiking in layer 5 (L5; Supp. Fig. 5a, b). Similar changes are observed in local field potential (LFP) spectrograms from different cortical layers (Supp. Fig. 5c). These results further confirm our hypothesis that inhibition of upper cortical layers generated disinhibition of neurons in lower cortical layers.

### Stimulation during behavior

To demonstrate our platform’s applicability for behavioral neuroscience experiments, we trained the monkeys to perform a center-out reach task using their right hand (Fig. 7a) collecting data over multiple sessions (e.g., Supp. Fig. 6). To begin, the monkey held his finger at a start target in the center of a screen, and then the end target displayed on the screen randomly in four locations: right, up, left, and down. We applied optical stimulation for 900 ms with all LEDs of our red 4 × 4 array (∼12 mW/mm^2^) simultaneously for a randomly selected 50% of the trials with the expectation that optogenetic inhibition would impede reach. The onset of the stimulation aligned with the appearance of the end target. After a random delay, an audial “go-tone” would play, signaling the monkey to reach for the end target. Monkeys typically completed the reach before the end of stimulation. Total counts of successful trials are reported in Supp. Table 1.

**Figure 7:**
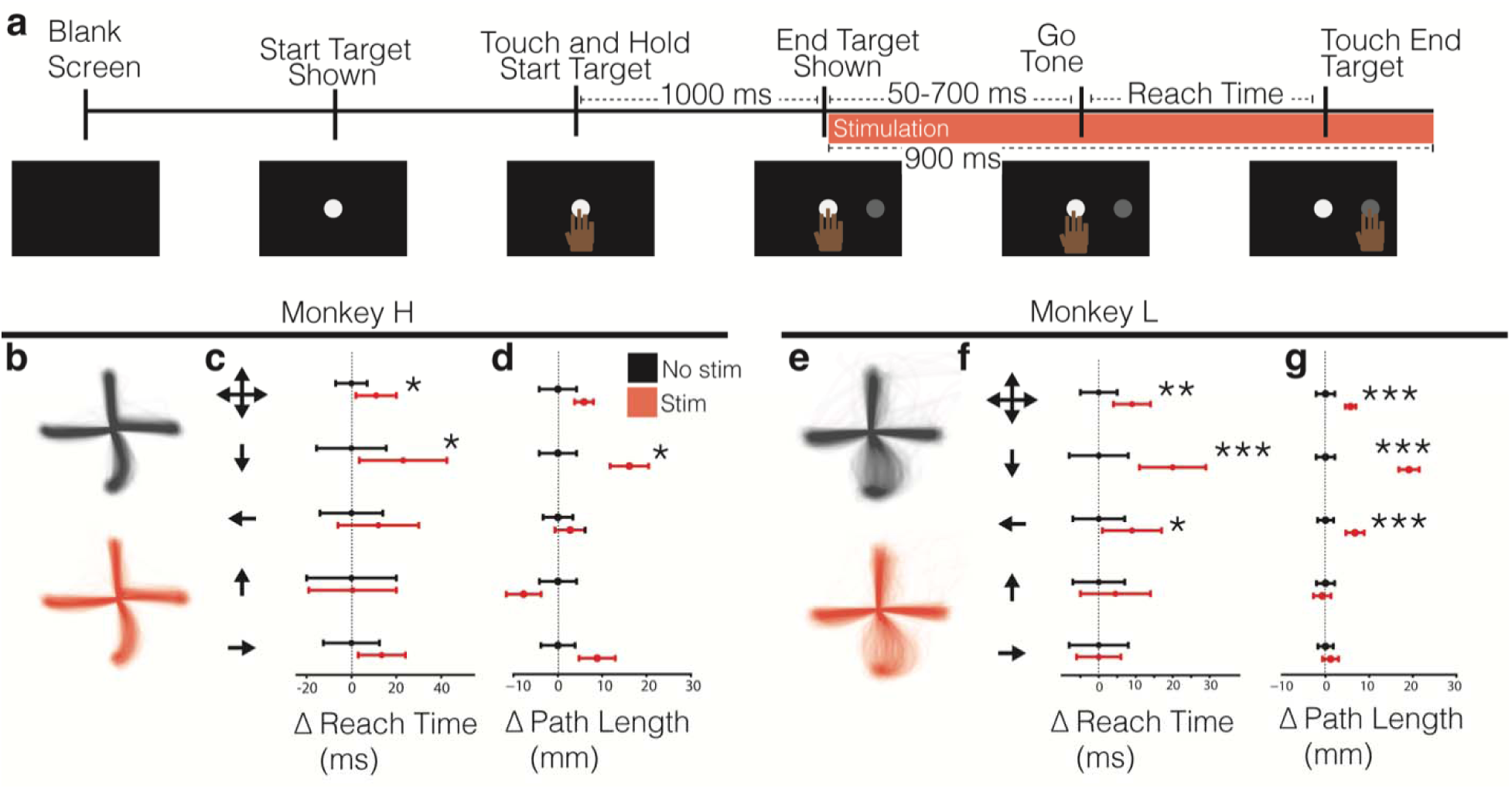
Behavioral task. **(a)** Schematic of behavioral task and stimulation timeline (neither to scale). The start and end targets were 25 mm diameter, and the center of the end target appeared 80 mm away from the center of the start target. **(b, e)** Reach trajectories in 2D. Differences in reach times **(c, f)** and path lengths **(d, g)** for Monkeys L and H, respectively, normalized to the non-stimulated median for each condition. Arrows on the left indicate reach, where all four arrows together (top row) represent the aggregation of reaches in all directions. Median and 95% confidence intervals are reported as error bars. Asterisks denote * p < 0.05, ** p < 0.01, *** p < 0.001 using the Mann-Whitney U test. See Supp. Table 1 for the number of trials.

Optogenetic inhibition led to significantly longer reach times in both monkeys (H: p = 0.04, L: p = 8.2e-3; Fig. 7c, f); however, trajectories of the reaches during stimulated and non-stimulated conditions do not look qualitatively different (Fig. 7 b, e). To better investigate these delays, we analyzed reach times for each movement direction individually (Fig. 7c, f). We found direction-specific effects in both animals. In Monkey L, reaches in the down and left directions were significantly impacted by stimulation (see Fig. 7f; direction-specific statistics in Supp. Table 2). In Monkey H, significant delays were observed in the downward direction as well (Fig. 7c, direction-specific statistics in Supp. Table 2). Additional analyses of path length were consistent with these findings (Fig 7d, g).

We analyzed the neural activity recorded during reaches with and without optogenetic stimulation to identify the change in neural activity underlying the reach disruption (see STAR Methods for details on the neural data analysis; data adapted from ^57^). We plotted heatmaps of theta-band activity (4-10 Hz) during rest and during the planning phase of the task just before go-tone (Fig. 8a, b). Theta band is known to be reflective of behaviorally relevant cortical activity. We found that area 7 exhibits increased activity during the planning phase. Next, we selected example electrodes in opsin expressing and deficient regions of area 7 and analyzed the signal during the planning period of the task with and without stimulation (Fig. 8c-e). To assess the changes in local activity of the selected electrodes, we analyzed high-gamma band activity. For both animals, both electrodes exhibited similar increases in baseline-normalized high-gamma-band activity during the task in the absence of stimulation. For Monkey H, both electrodes had statistically significant increases in activity when stimulated (expressing: p = 9.0e-31; deficient: p = 2.1e-12; Fig. 8f). For Monkey L, only the activity of the electrode in the opsin expressing area had increases to a statistically significant degree (expressing: p = 3.8e-74; deficient: p = 0.09; Fig. 8g). Regardless, for both monkeys we found that the stimulation-evoked activity of the electrode over the opsin expressing area was more pronounced than the electrode over the opsin deficient area to a statistically significant degree (Monkey H: p = 2.0e-7; Monkey L: p = 5.7e-35), which was not the case in the absence of stimulation (Monkey H: p = 0.39; Monkey L: p = 0.08; Fig. 8f, g). These results support our finding that optical stimulation of neural tissue expressing Jaws increased ECoG power both during behavior and at rest (Fig. 6).

**Figure 8:**
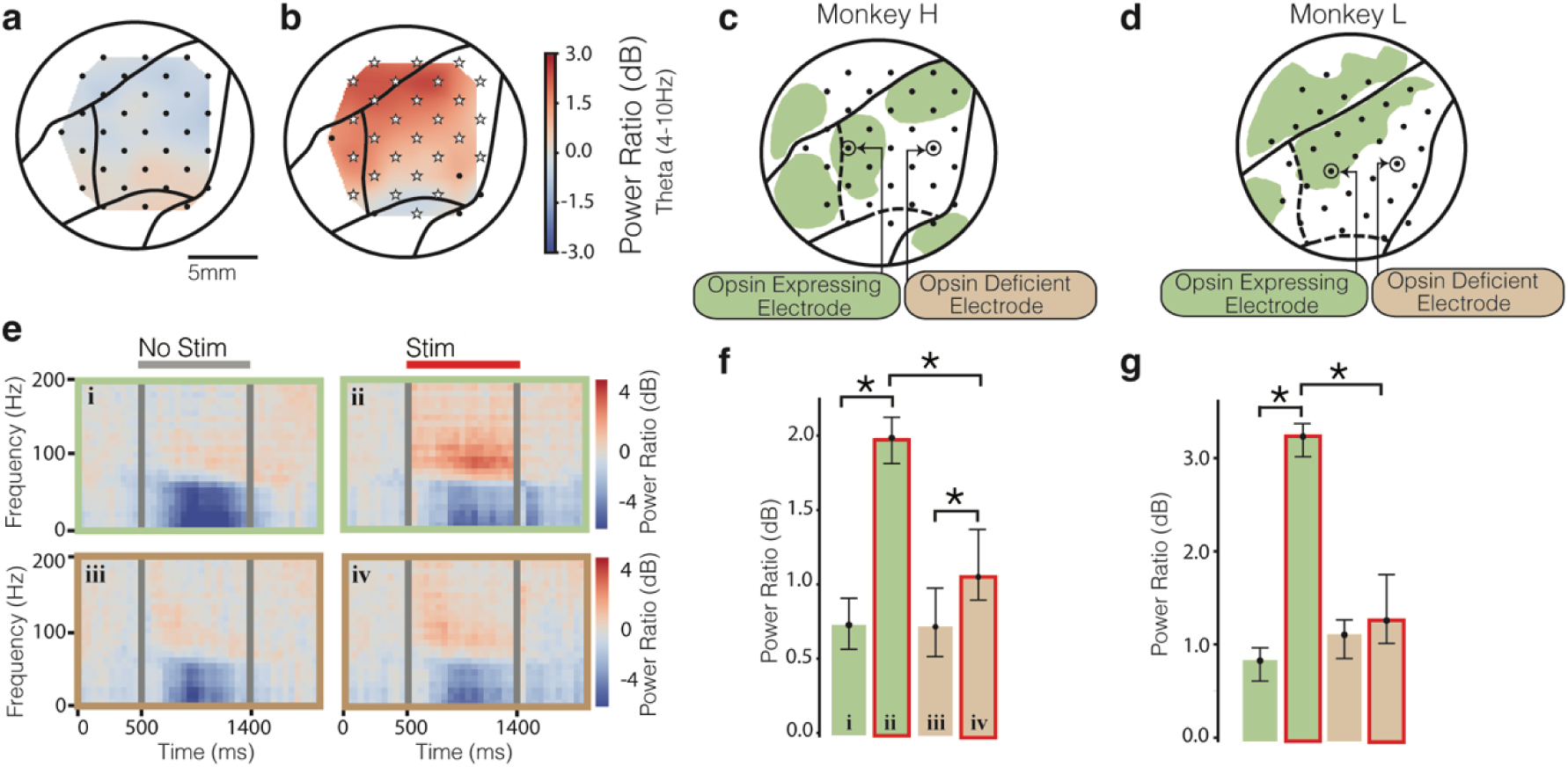
Effect of opsin expression status on neural activity during reach. Theta-band (4-10 Hz) ECoG heatmap of resting **(a)** and movement planning (**b**) activity in the absence of stimulation for Monkey H. White stars on electrodes indicate statistical significance (p < 0.05, one-sample t-test). (**c, d**) Epifluorescence map for Monkey H (c) and Monkey L (d) indicating electrodes selected for use in (e, f) and (g), respectively. **(e)** Trial average spectrograms from individual electrodes called out in (c) during reach. (i): opsin expressing electrode, no stimulation. (ii): opsin expressing electrode, stimulation. (iii): opsin deficient electrode, no stimulation. (iv): opsin deficient electrode, stimulation. Spectrograms are average neural activity across trials for the 900 ms stimulation (or where stimulation would have been) and the 500 ms before the onset of the stimulation and 900 ms after the offset of stimulation. Grey bars indicate bins during the first 50 ms of stimulation onset and offset – data during these times are excluded due to residual artifact after artifact removal. Spectrograms were averaged over 1 electrode and 253 trials in the case of no stimulation, and 241 trials in the case of stimulation. **(f, g)** Median high-gamma-band (80-200 Hz) ECoG power during the task (900 ms) for the aforementioned electrodes with and without stimulation for Monkey H (f) and Monkey L (g). For Monkey H, trials are the same as those reported in (e). For Monkey L, 440 trials with stimulation and 401 trials without stimulation were used. Whiskers are 95% confidence intervals of the median generated with a bootstrapping method, and asterisks indicate significant differences between the conditions (Bonferroni correction applied to p < 0.05).

## Discussion

We present an NHP optogenetic ECoG platform both accessible to researchers with diverse expertise and flexible for diverse experimental aims. Our work entails five complimentary technological advancements tied together with the common thread of experimental flexibility and researcher accessibly: an adaptable and modular platform, a chronic MMAD, scalable optical hardware for patterned stimulation, improved ECoG-compatible optical access, and efficient viral vector delivery without live-MRI monitoring. We used optical imaging, electrophysiology, and behavioral data to showcase our platform.

We designed our platform to be widely accessible by making it modular and replicable in several ways. All parts are interchangeable between monkeys except those chronically implanted. Our MMAD molding process did not require any special facilities or tools aside from the custom mold pieces, and the miniature ECoG array is commercially available (Ripple Neuro Inc.). We have previously demonstrated the MMAD’s capability of electrical stimulation ^41,58–61^, which opens further experimental opportunities, especially given that optogenetic and electrical stimulation have been shown to work synergistically ^62,63^. Furthermore, we predict our work will spur the development of MMADs with higher channel counts of smaller and transparent electrodes to increase electrophysiological resolution and optical access ^64,65^. In particular, we expect future versions of the MMAD with electrode sizes in the order of 10-30 µm to be capable of recording action potentials or multiunits from the cortical surface, similar to previous work ^64,66,67^. We also expect LEDs will be incorporated into future MMAD designs, much like past work with electrodes arrays (e.g., ^68,69^). Our platform can facilitate iterative development in MMAD design because explantation of past designs and re-implantation of future designs is minimally invasive. Replacing or updating electrophysiological or optical stimulation hardware implanted with traditional window-less approaches, like Utah array-style devices ^70^ or LED arrays sutured to the native dura ^34^, is more surgically complex in comparison to our window-based method.

The ring and clamp connector tray facilitate experimental flexibility for imaging, electrophysiological recording, and both fiberoptic- and LED-based optical stimulation. Most of the optical stimulation components can be 3D printed in-house with common 3D printers, which facilitates quick and inexpensive revisions for experimental needs. Parts can be soldered onto the LED-related PCBs in-house with minimal experience, and simple, custom-designed battery-powered circuitry mitigates line-noise. The LEDs are commercially available in many colors and have the same footprint, which facilitates custom LED arrangements including multi-color arrays for step-function opsins ^71^. Our custom circuitry does not use pulse-width modulation, but drives the LEDs continuously. Continuous stimulation minimizes the count of optical stimulation onsets and offsets per stimulus, thus mitigating complexity of photo-induced artifacts in electrophysiology data. Regardless, the remaining singular onset and offset of continuous stimulation produced artifacts in our ECoG data. We rejected the artifacts with a rigorous procedure designed to preserve the neural responses of stimulation. Our artifact rejection method can be readily adapted for other experiments combining optogenetics with electrophysiology. We incorporated an air gap into our LED setup to mitigate tissue heating, which is a concern in all optogenetic experiments, and particularly when placing LEDs on ^34,35^ or in the brain. To further protect the brain from heating, we designed the LED array and the driving circuitry to be separate PCBs to keep heat produced by the driving circuitry (i.e., opamps and resistors) away from the brain. Our temperature measurement data (Supp. Fig. 2) indicate that our designs were effective at limiting the cortical temperature change to less than 1 ℃ during our stimulation protocols (Supp. Fig. 2), which is below the safe limits for the brain ^44–47^ and has minimal impact on neuronal activity ^48,49^. The stimulation setup of previous work allowed for the optical stimulation of only two or three point-locations at a time due to the physical interference of fiberoptic wires within the chamber ^25^; in contrast, our LED array provides large-scale patterned optical stimulation for targeting several brain areas simultaneously. Additionally, our platform allows users to monitor both stimulation and recording locations every experimental session for displacement with respect to the brain (ref. Fig. 3a), which provides a great advantage over surgically implanted devices with limited visibility of the brain (e.g., ^20,34,35^). Similarly, given that our platform can be integrated with optical techniques such as optical coherence tomography angiography (OCTA) ^41^, calcium imaging and intrinsic signal imaging, it is well-suited for studying neural and vascular dynamics—including neurovascular coupling—across timescales ranging from minutes to months, and in conjunction with behavioral tasks. We anticipate development of higher-count and multicolor LED arrays to further complement our large-scale optical platform, which would be especially powerful with multi-opsin and step-function opsin experiments ^16^. The design files of our electronic and hardware parts are publicly available (see Data and Code Availability).

Our work pushes the bounds of previous efforts across the field largely hindered by the challenge of maintaining optical access and especially at the large scales that will be necessary to study complex network interactions across multiple brain regions in awake behaving NHPs. As with past work, we resected both the skull and the opaque native dura and implanted transparent artificial duras that are biocompatible and can provide optical access to the brain for months ^23,25,72^ before tissue growth stifles optical access ^25^. This tissue is a neo-membrane and cannot be easily removed during early stages of growth because it is nourished through the brain, but the tissue can be surgically removed once the neo-membrane matures and separates from the brain. This growth process forces the researchers to wait for about one to two months before resuming experimentation, and thus highlights the importance of maintaining optical windows for an order of months, as others have done with classic optical windows and as we have now demonstrated with an electrocorticographic optical window.

Early works attempting to integrate electrode arrays and large-scale optical windows did not result in long-term optical access ^25,29^. In this work, we achieve over 3 months of optical and electrophysiological access by embedding electrodes into a transparent polymer ^41^ and then integrating the array into an artificial dura to create a single MMAD. The brain’s response is similar to classic, electrode-free silicone artificial duras. While others have also embedded miniature ECoG arrays into artificial duras ^73^, their ECoG design restricted optical access. We integrate our MMAD, purposefully designed for optical access ^41^, with an artificial dura to support chronic optogenetics, imaging, and ECoG.

While others have proposed window-less designs for LED-based NHP optogenetics ^20,34,35^, our work facilitates routine epifluorescent monitoring of large-scale optogenetic expression, revealing experimental insights otherwise challenging to ascertain. For example, in our experiment the optical window empowered us to confirm that CED performed without the complexity of live-MRI (unlike ^25^) resulted in a monitoring progression of large-scale expression up to a plateau in expression at 5 weeks. The large areas of epifluorescent coverage are comparable with previous work ^25^. Around 2 years later we observed a waning of epifluorescent expression, which again highlights the value of window-based designs for routine optical monitoring of the cortex. The viral reinjection and subsequent epifluorescent confirmation in Monkey L further underscore the importance of the optical access enabled by our setup. Our platform also supports non-genetic optical techniques, such as infrared neural stimulation ^26^ and photochemical lesioning ^28,58,59,74,75^, which sets the stage for such studies to be complimented with chronic ECoG data via our MMAD. Further, our platform enables routine monitoring of tissue growth, which can be problematic for many optical techniques and is difficult to assess without an optical window. Granted, our platform does not provide optical access beyond the cortical surface, yet future iterations of our platform could be combined with hardware for depth recording and stimulation.

Our CED success is an important proof of concept for our method of optogenetic CED in monkey cortex without live-MRI guidance. While live-MRI is a powerful method of visualizing infusions in real-time during surgery ^25,50^, the method is logistically challenging or infeasible at many institutions, limiting its use. Our MRI-free CED method was performed entirely in the surgery suite and during the same surgery where all other surgical operations were performed. This eliminated the need for MRI-compatible syringe pumps and complicated fluid lines, and instead used a setup which may be widely accommodated by surgery suites at other institutions, while still capitalizing on the efficiency of CED over traditional diffusion-based approaches. While with this technique we lose the on-line monitoring of infusions, our previous work provides a comprehensive bench-side technique to model the CED infusion profile prior to surgery ^76,77^. This bench-side model was informed by the correlation between CED parameters and on-line MRI monitoring of infusion profiles in NHP brains. This modeling technique is predicated on a reflux-free infusion, which our new CED technique empowered us to monitor. Together, the model and method support accessible and comprehensive planning and execution of widespread viral delivery in NHP brain.

Based on previous studies^9,25,53^, we expected that the strongest effects of optogenetic stimulation would occur in the opsin expressing areas and that weaker effects would occur in opsin deficient areas, and our data support this. The weaker effects observed in opsin-deficient areas (as seen in Monkey H, Fig. 8f) may be attributed to one or more of the following factors: (1) opsin expression that is too weak to be detected via epifluorescent imaging; (2) propagation of neural responses originating in opsin-expressing regions, leading to secondary effects in opsin-deficient areas through network-level interactions; and (3) potential thermal effects in opsin-deficient regions. However, our bench-side temperature measurements indicate that stimulation-induced temperature changes are less than 1°C, suggesting that the likelihood of thermally induced neural activity is minimal.

While the relative strengths of stimulation-evoked responses with respect to opsin expression were aligned with our expectations, the nature of the responses were surprising. We originally hypothesized that our inhibitory opsin would reduce ECoG signal power when stimulated, but found a general increase in ECoG signal power regardless of behavioral state (Fig. 5, 6, 8), including when channels were selected or grouped by the presence of expression in their vicinity (Fig. 6, 8). To explain this finding, we speculated that upper layers of the cortex, which are known to have higher ratios of inhibitory neurons than other layers ^78,79^, were more strongly inhibited due to greater proximity to the light source, and thus the electrodes netted higher electrophysiological signal power from the lower cortical layers that received less inhibition from the higher cortical layers. In particular, layer 2/3 is known to project significantly to layer 5, which is a cortical layer that contributes heavily to EEG ^80^ and µECoG ^81^ recordings. Indeed, we have previously shown with electrical stimulation in rat brain that lower rates of neuronal firing in layers 1/2 are correlated with higher rates of firing in layers 5/6 (and vice-versa) ^82^, and that ECoG activity is more strongly correlated with the spiking and LFP activity of layer 5/6 than other cortical layers ^83^. This reasoning agrees with our in vivo findings, and to further investigate we have performed a simulation study in which we observed that illuminating Jaws-expressing cortex caused inhibition of upper cortical layers and excitation (disinhibition) of lower cortical layers. These results further validated our in vivo ECoG responses.

Simultaneous depth and surface recording has shown that an increase in firing in lower cortical layers is correlated with an increase in high-gamma ECoG power ^49,83^. However, in this work we observed not only an increase in high-gamma power but also in other frequencies. This finding is aligned with other work that showed frequency-specific responses to different stimulation parameters ^84^. Our light-evoked changes in ECoG signal across different frequencies could also vary depending on stimulation parameters, which merits additional study.

Although increase in signal power during optogenetic inhibition was unexpected, our control experiments validated that the increase in signal power was, indeed, optogenetically evoked (Figs. 5, 6). Lasers generated more spatially specific modulation than LEDs, and red light provided stronger neural modulation than blue light in accordance with the spectral characteristics of Jaws ^52,53^. The latter observation would not be expected had neural responses been merely an effect of tissue heating as blue light is absorbed more than red light and thus causes more spatially specific heating ^85^. Furthermore, as explained above our bench-side temperature measurements suggest that the likelihood of thermally induced neural activity is minimal. In addition, when the channels were grouped by the presence or deficiency of expression in their vicinity, red stimulation through tissue growth and blue stimulation with clear optical access experiments failed to generate statistically significant results, leaving red stimulation with clear optical access, i.e., both the optimal wavelength and optical access, as the only case where opsin expressing areas were evoked to higher power than opsin deficient areas (Fig. 6).

Past studies have attempted to use excitatory opsins to evoke musculoskeletal activity in NHPs at rest. Early trials were unsuccessful ^16^, but more recently, a proof-of-concept was demonstrated ^86^. A preprint study has shown disrupted sensory-guided reaching behavior using excitatory optogenetics ^87^. Other studies have used inhibitory opsins to produce behavioral effects; Jaws has been used in rhesus macaques to produce behavioral effect in saccade tasks ^53,88,89^ and a joystick release task ^89^. To demonstrate the versatility of our platform for behavioral studies, we applied optogenetic stimulation of Jaws expressing areas in the PPC during a center-out reach task, hypothesizing that this deactivation would result in delayed reaching movements. This hypothesis is based on the relationship between the PPC and motor intentions ^90–103^, and neuronal connections between the PPC (including areas 5 and 7) and the motor and premotor cortices ^104–109^. PPC acts as a sensory integration center, downstream of pre-motor regions. Previous studies have inhibited the PPC through cooling and found reaching movements were delayed ^110^. Additionally, single unit recordings in the PPC verified the proactive role of this brain region in motor planning during a center-out reaching task ^111^. While future PPC studies could use our platform to disentangle interrelated PPC functions, here we present an alternative to current methods that allows for precise probing of the region with optogenetic deactivation.

Our behavioral effect is comparable with other work: For example, electrical subthreshold microstimulation of rhesus macaque premotor cortex, an area shown to be directly responsible for preparation of motor movement, impeded reaction time of a reach task by 10s of ms ^112^. In both of our monkeys, we attain similar results by optogenetically stimulating the PPC, an area indirectly responsible for preparation of motor movement ^113^. Our results indicate that our platform is suitable for studying neural mechanisms of behavior with millisecond-precision, spatial specificity, and reversable modulation, with behavioral effects comparable to prior electrical stimulation work.

Our results support our hypothesis that optogenetic inhibition disrupts reaching behavior; however, reach time was impacted differently in our two monkeys, which may be due to different expression profiles over the PPC. Monkey L had larger behavioral perturbations in both reach path and reach time while also having greater opsin expression near the interparietal sulcus. This region of the PPC is known to have significant task related activity ^114^, which agrees with our results and may indicate that we perturbed task-related activity more precisely in this monkey. This observation is enabled by our ability to chronically monitor opsin expression and its overlap with neural activity. To our knowledge, this study is the first report of reach disruption using an inhibitory opsin in NHPs. Further, our study uses a brain area not directly responsible for motor actions, as opposed to the motor and premotor cortices more popular in musculoskeletal behavioral experiments, which highlights the range of experiments our platform can facilitate.

Collectively, our methods make cutting-edge neuroengineering techniques accessible to NHP researchers with diverse expertise. Our results validate our chronic setup and indicate its suitability for large-scale and long-term stimulation and recording experiments in behaving monkeys. The combination of high temporal and spatial precision with large-scale and long-term access renders our platform particularly useful for experiments which need fine-scale manipulation and sensing over longer timescales and larger brain areas. Examples include studies of network-level activity, functional connectivity between different cortical areas ^9,115,116^ and neurological diseases and disorders.

Given that not every experiment requires every technology or use-case we present, our platform is functionally modular, providing flexibility to select modalities appropriate for one experiment while maintaining the opportunity to incorporate other modalities in future studies with the same monkey. For example, an imaging-specific neuropathology study could be followed by development of simultaneous optogenetic and electrical stimulation protocols, which could speed the translation of the new diagnostic or therapeutic techniques from animals to humans. We anticipate wireless closed-loop technologies ^117^ and next-generation optical and ECoG devices ^118,119^ will be integrated into designs based on our platform to make our work practical for experimentation on freely moving animals. We expect this to further advance stimulation-based therapies for neurological diseases and disorders ^120^.

## Acknowledgments

We thank Zachary Ip and William Ojemann for their help with MRI processing; William Ojemann, Kali Coubrough, Jia Wen Chan, Zada Anderson, Stephen Philips, and Andrew Graham Johnson for their help with part design; Ripple Neuro, especially Alex Johnson, Jose Ortega, and Jessi Michel, for their help with designing and manufacturing the MMADs; William Ojemann for his help with MMAD molding; Marcus Chu and Mona Rahimi for their help with electronics; Kelly Morrisroe, Tiphaine Belloir, Kelly Yeh, Toni Haun, Briana Smith, Kali Coubrough, Shivalika Chavan, Pavi Rajeswaran, Kiara Evitts, and Avi Matarasso for their help with animal training; Jane Sullivan for her help with opsin validation prior to infusion; Greg Horwitz, Keith Vogel, Toni Haun, Zachary Ip, and WaNPRC veterinarians and staff for their help with surgeries; Tiphaine Belloir, Kelly Yeh, Toni Haun, and WaNPRC veterinarians and staff for their help with chamber care; Kali Coubrough and Shivalika Chavan for their help with behavioral rig development; Felix Schwock and Sophia Shan for their help with data collection; Ryan Ressmeyer and Arie Aelmore for their help with data analysis; Anne Pierce for her guidance with VERTEX usage; and our monkeys for their invaluable participation.

We thank our funding sources: R01 NS119395 (DJG, JB, JZ, KK, LS, EEF, AY), R01 NS116464-01 (AY), University of Washington Big Data for Genomics & Neuroscience Training Grant (JB), Weill Neurohub (SF), Washington National Primate Research Center P51 OD010425 (DJG, JB, NS, JZ, SF, AY), American Heart Association (AY), 1T32 MH132518 (NS), R01 MH125429 (NS, AY), K01 AG071798 (HJ), D-SPAN F99NS130828 (KK), and the National Science Foundation Graduate Research Fellowship Program (KK). DJG is currently funded by K25 AG086663.

## Author contributions

Author CRediT: AY conceptualized the project, provided supervision, and acquired funding; DJG, JB, KK, SF, LS, EEF, HJ, and AY developed the methodology; DJG, JB, JZ, KK, and LS performed the investigation; NS, JB, LS, and HJ developed and implemented the software; DJG, NS, and JB performed the data curation and formal analysis; DJG, NS, JB, JZ, SF, and LS performed the visualization; DJG wrote the original manuscript; all authors reviewed and edited the manuscript.

## Data and code availability

Our data will be made available upon reasonable request. Our analysis code for in vivo data and the design files of our electronic and hardware parts are publicly available (https://bitbucket.org/yazdanlab/interface_code/src/main/; https://bitbucket.org/yazdanlab/interface_electronics/src/main/; https://bitbucket.org/yazdanlab/interface_hardware/src/main/).

## STAR Methods

### Multi-modal artificial dura (MMAD)

We embeded a multi-modal artificial dura (ECoG array), similar to past work ^41^, into a traditional artificial dura (Fig. 1b). Briefly, the MMAD consisted of four conductive layers sandwiched between five transparent layers of medical-grade copolymer, where the conductive layers were aligned with each other to maximize optical access. Conductive layers were composed of platinum particles dispersed in a matrix within the polymer for mechanical flexibility. This design eliminated the need for attached printed circuit boards (PCBs), enabling long-term implantation without the challenges of protecting PCBs from moisture and fluids within the chamber. The MMAD was developed in close collaboration with Ripple Neuro (Salt Lake City, UT, USA), who manufactured the arrays. The company’s sales team may be contacted for ordering details.

A traditional artificial dura is typically described as “top-hat” shaped: a single silicone piece is composed of a cylindrical wall with an optical window at the lower end and a flange extending under the native dura to reduce regrowth of the native dura (Fig. 1b) ^23–25^. Here, we embedded the MMAD into the artificial dura with a simple, previously described molding process ^40,42^. The molded device is an artificial dura (skirt Ø: 31.2 mm; skirt thickness: 0.4 mm; wall outer Ø: 21.0 mm; wall inner Ø: 18.6 mm; height: 11.9 mm) with a fully embedded MMAD to allow for electrical recording and stimulation from 32 electrodes (Fig. 1b).

We designed the entire MMAD, including the cables, as a corrosion resistant single piece. We housed the cables of the MMAD in the chamber between experiments, and during experiments we attached the cables to the electrophysiological recording and stimulation hardware.

### Chamber

The chamber is a cylinder (internal Ø 23 mm), 3D-printed and milled from titanium (Ti-6Al-4V, Hybex Innovations, Anjou, Quebec, Canada), which provides a sturdy base for experimental equipment and protects the brain when the monkey is freely moving between experiments, similar to previous work ^25^. To design the chamber to fit the skull of each monkey, we developed a software pipeline (MATLAB, MathWorks, Natick, MA, USA; Solidworks, Waltham, MA, USA) to extract the skull curvature from an MRI file and then used the curvature to design a chamber with a unique skull-fitting skirt (Fig. 1a, c; Supp. Fig. 7) ^39,40^. We included an intra-cranial rim under the skirt (Supp. Fig. 7) to aid in positioning the chamber on the skull during surgery and to reduce bone growth around the wall of the craniotomy, and included 12 holes in the skirt (Supp. Fig. 7) for titanium skull tap screws (Crist Instrument Company, Hagerstown, MD, USA). We incorporated concentric tracks within the chamber to store the MMAD cables between experiments (Fig. 1b-d). We threaded the chamber with a high thread pitch (M38 × 1.5) to allow either affixation of experimental equipment or a cap to close the chamber between experiments (Fig. 1c; Supp. Fig. 7).

### Cap

We designed a cap (stainless steel 304, machined by Hybex Innovations) to close off the chamber between experiments. We applied polytetrafluoroethylene (PTFE) tape to the threads before each closure to prevent binding of the threads. We used a set screw (HSS11400187HD, #4-40 × 3/16) to fasten the cap shut against the top rim of the chamber.

### Electrophysiology hardware

Our electrophysiology setup included a ring, a clamp connector tray, and PCB clamp connectors (Fig. 1e, f). To prepare our platform for recording and stimulation, we screwed the ring onto the chamber and secured the clamp connector tray onto the ring with 2-4 screws (M2.5 × 6). The ring was machined from stainless steel (304; Hybex Innovations) and the clamp connector tray was 3D printed with titanium (TI-64; i.materialise, Leuven, Belgium).

The ring had 32 screw holes (M2.5) around its rim to offer optimal flexibility in the orientation of attachments. The upper face of the ring included a slight recess just above the threads for additional hardware to be easily aligned with the ring for fastening.

The bottom of the clamp connector tray included an intra-ring rim around the window to fit into the recess of the ring. The base of the tray included a right-angled outer rim for grounding electronics as well as four elevated screw holes (M2.5 or 4–40 thread) designed for optical stimulation equipment to be secured to the tray. We designed the tray to be compatible with both this work’s LED-based optical stimulation setup and our lab’s previously described fiberoptic-based setup ^25^. We designed the arms of the tray to be low weight and provide support for the MMAD cables as they extended from the artificial dura walls at 45°. The distal end of the tray arm was approximately 9.5 cm away from the cranial window.

The arms of the tray supported PCB clamp connectors (Fig. 1e, f; Ripple Neuro) held in place by rubber bands (not shown), which secured the MMAD cables approximately 4.5 cm away from the brain and provided an electrical connection between the cables and commercial neurophysiology equipment (Grapevine Nomad, Ripple Neuro). Two of these setups may be simultaneously deployed bilaterally, although this work is exclusive to the left hemisphere.

### Optical stimulation hardware

To optically stimulate across large cortical areas (∼1 cm^2^), we incorporated either a 4 × 4 or a 3 × 5 LED array (Fig. 2). The full 4 × 4 array was used for stimulation during reaches. To prevent LED arrays from moving when in use, we designed a system involving a rod, tube, coverslip, and stimulation holder (Fig. 1g). We custom designed and 3D printed the rod, tube, and stimulation holder in-house with FDM PLA (M2 revG, MakerGear LLC, Beachwood, OH, USA). We glued (Better Ultimate Adhesive, slow dry, Cemedine Co. Ltd., Shinagawa City, Tokyo, Japan) the coverslip (Ø 16 mm, ORSAtec, Bobingen, Germany) to one end of the tube. The base, which we affixed to the clamp connector tray, secured the coverslip against the top side of the MMAD and a set screw (HSS11400187HD, #4–40 × 3/16) secured the rod inside the tube. The rod positioned the LED array over the brain and above the coverslip. Our setup created an air gap between the LEDs and the coverslip, which protected the brain from conductive heating. Four screws in the base secure the tube in place.

The LED array consisted of a circular custom rigid printed circuit board (PCB; Ø 15.5 mm; OSH Park, Portland, OR, USA). The LEDs (L1C1-RED1 or L1C1-BLU1, Lumileds, Schipol, Netherlands) of the 3 × 5 array were individually addressable, while the 4 × 4 array was wired to drive four individually addressable columns of LEDs. Both PCB versions included an internal copper plane for heat dissipation. We soldered the LEDs on one side of the PCB and a connector (501331-0807, Molex Inc., Lisle, IL, USA) on the other side. We drove the LEDs with a custom external PCB (OSH Park) connected to the LED array by a cable assembly (151330806, Molex Inc.). The external PCB consisted of voltage-controlled current sources. Each driving circuit consisted of a high output current opamp (TLE2301, Texas Instruments, Dallas, TX, USA) with a capacitor between the compensation network terminals (not shown) and a resistor (15 Ω, 5 W, SMW515RJT, TE Connectivity, Schaffhausen, Switzerland), where the resistor value controls the ratio of input voltage to output current. The input voltage is provided by our neurophysiology system’s analog voltage output (Ripple Neuro) and is prevented from floating high (e.g., when the neurophysiology system is turned off) by a pull-down resistor (10 kΩ). We powered our PCBs with two in-series batteries (9V Li-ion rechargeable, Keenstone, Industry, CA, USA) to reduce line noise in our electrophysiological recordings. We depict our circuit and arrays in Fig. 2 and Supp. Fig. 1.

We also designed our clamp connector tray to be compatible with our previous laser stimulation setup ^29^, and its slightly modified version is briefly described here. Our setup was composed of a base similar to the base for the LED array setup, and a stimulation ring, both FDM 3D-printed in-house with black PLA. The base was screwed to the clamp connector tray. The stimulation ring guided telescoping cannulas (HTX-13R, HTX-16T, and HTX-18T, Component Supply Co., Sparta, TN, USA) to different stimulation locations, and optical fibers were slid through the cannulas. We coupled lasers (520 nm or 638 nm, LDFLS_520_520_638_638, Doric Lenses, Quebec, Canada) to the optical fibers.

### Animals

We recruited two healthy, socially housed male rhesus macaques (Macaca mulatta; Monkey H: 9 y, 13 kg; Monkey L: 9 y, 11 kg) for this study. Water was available ad libitum and standard food rations were provided after experimental sessions. All animal care and experiments were approved by the University of Washington’s Office of Animal Welfare, the University of Washington’s Institutional Animal Care and Use Committee, and the Washington National Primate Research Center.

### Surgical procedures

We performed surgical procedures on both monkeys similar to procedures in other work ^25^. We sedated the monkey and shaved its head before anesthetizing and head-fixing him in a stereotactic frame. The respiratory rate, heart rate, and body temperature of the monkey were monitored throughout the procedure. We created an incision several cm long to the left of the head’s midline and used elevators to peel back the skin, musculature, and muscle fascia. We confirmed the location of the center of the craniotomy with stereotactic coordinates. Using a Ø 25-mm trephine, we created the craniotomy. Then we resected the dura by lifting the dura with a curved needle and trimming with ophthalmic scissors, revealing the posterior parietal cortex. After test-fitting the titanium chamber to the skull, we placed a flat transparent silicone artificial dura on the brain to keep the brain moist.

With the craniotomy complete, we performed convection-enhanced delivery (CED) of Jaws (rAAV8/hSyn-Jaws-KGC-GFP-ER2, 5.4 x 10^12^ genome copies per milliliter (gc/mL), University of North Carolina Vector Core), a red-shifted inhibitory viral vector ^52^. Prior to surgery, we had manufactured a 1-mm stepped-tip silica cannula (inner cannula: inner Ø 320 µm, outer Ø 435 µm; outer cannula: inner Ø 450 µm, outer Ø 673 µm; Polymicro Technologies, Phoenix, AZ, USA) for CED similar to our previous work ^25,51^ by applying cyanoacrylate (super glue) to the inner cannula, sliding the outer cannula over the inner cannula, and using a razor blade to trim the inner cannula extending from one side of the outer cannula to make a 1-mm stepped-tip. We removed the artificial dura from the brain, punched a hole through the artificial dura to be used for our injection, laid the artificial dura back on the brain to keep the brain moist, and then inserted the cannula tip through the hole approximately 2 mm into the cortex (Fig. 3a, left) and began the infusion protocol using convection-enhanced delivery similar to previous work ^25^. We infused up to 50 µL at each of several locations across the cortex. Once complete, we placed a new artificial dura on the brain.

Unlike previous work where live-MRI was performed during infusions ^25,50^, we performed the infusions without live-MRI. This allowed CED to be performed in the surgery suite while we were able to visually inspect the location of infusion for any reflux. It is also eliminated the need for MRI-compatible syringe pumps or complicated fluid line setups – the syringe and cannula were attached directly to the pump, and a stereotactic arm was used for cannula insertion during CED.

Following CED, we used a piezo-drill (5120063AS animal science package, insert tips IM1-AL and IM1S, Piezosurgery Inc., Columbus, OH) to drill screw holes and used titanium tap screws (Crist Instrument Company) to secure the chamber to the skull. We removed the artificial dura and placed the MMAD on the brain and housed the cables in the slots of the chamber (Fig. 1d). Finally, we applied PTFE tape to the threads, closed the cap, secured the set screw, and applied bone wax to the set screw. Having secured and closed the chamber, we closed the wound using standard surgical techniques.

We had implanted a titanium headpost (Crist Instrument Company) on each monkey prior to this study^39^.

### Chamber cleaning and hardware assembly

We developed a process of cleaning the chamber at least three times each week, starting one week after the surgery. We briefly describe the processes here. After head-fixing the monkey, we cleaned the cap, chamber margin, surrounding scalp and outside of the ear. Once clean, we removed the cap and cleaned the margin. We used sterile cotton-tipped applicators dipped in saline to clean the threads and the rim of the chamber. To clean the inside of the chamber, we used a sterile syringe and blunt-tipped needle to irrigate warm sterile saline on and in the MMAD, its cables, and the storage area for the cables.

To prepare for electrophysiological recording, we screwed a sterile ring onto the chamber and screwed a sterile clamp connector tray to the ring with sterile screws. Then we removed the cables from the chamber and placed them into the clamp connectors, and rubber banded the clamp connectors to the clamp connector tray arms.

To prepare for optical stimulation, we attached the stimulation holder for either the LED array or for the lasers. In the case of laser stimulation, we slid the stimulation ring down around the stimulation holder and inserted the fiberoptic cable through the cannula and placed the cannula on top of the brain through the stimulation ring. In the case of LED stimulation, we assembled the LED array, rod, and tube with coverslip, and secured them together with a set screw. Then we lowered the assembly down through the stimulation holder and into the MMAD, and secured the tube into place with set screws. Black foil was wrapped around the stimulation equipment to prevent the monkey from seeing any lighting changes in the experimental rig due to stimulation.

When the experiment was complete, we removed the stimulation and recording equipment, cleaned the MMAD, and used sterile forceps to return the cables back into the chamber. Then we cleaned the chamber using sterile saline irrigation and applied antibiotic to the center of the MMAD. To avoid building antibiotic resistance, we rotated through polymyxin B sulfate, gentamicin, and amikacin sulfate antibiotics, applying one for 4-5 weeks and then switching to the next. To prevent the cap from binding to the chamber between cleanings, we wrapped a strip of sterile PTFE around a portion of the threads. Then we screwed on a sterile cap and a set screw.

### Epifluorescence imaging

We performed epifluorescence imaging of green fluorescence protein (GFP) by using a blue light for the excitation wavelength of GFP (NIGHTSEA stereo microscope fluorescence adapter, SFA-RB royal blue, Electron Microscopy Sciences, Hatfield, PA) in combination with a DLSR camera (Nikon D5300, Minato City, Tokyo, Japan) and a lens based on the emission wavelength of the GFP (Nikon DX AF-S NIKKOR 35mm 1:1.8G, SWM Aspherical ∞-0.3m/0.98ft Ø52) for imaging. We illuminated the brain from multiple angles to fully cover the optical window. The camera was fixed throughout the illumination process, which eased stitching the photos together with image editing software.

Expression coverage was measured by outlining the extent of expression in the stitched epifluorescent image manually, counting the number of pixels within the outlined areas, and converting to mm^2^ based on known distances in the image, i.e., the distance between electrodes.

### Behavioral task

Our center-out reach task (Fig. 7a) was controlled by a custom software in MATLAB using Psychophysics Toolbox version 3 ^121–123^, similar to previous work ^124^. A reflective sticker, placed on the middle finger of the monkey’s right hand, contralateral to the implant, was tracked by a motion capture system (Motive, OptiTrack, NaturalPoint Inc., Corvallis, OR, USA). These tracking data were processed in real-time by MATLAB. The monkey was rewarded for successful trials with juice (5-RLD-D1 D.A.R.I.S., Crist Instrument Company). The monkey was head-fixed and seated in a monkey chair inside a dark experimental rig. The visual and audio stimuli were provided by a monitor with built-in speakers within reaching distance of the monkey.

Monkeys performed several hundred trials each session (Monkey H: mean trials per session: 557.7, standard deviation: 217.6; Monkey L: mean trials per session: 715.7, standard deviation: 266.0). For Monkey H, there were 4,522 analyzed reach trials split over 8 sessions, while for Monkey L there were 11,101 trials split over 13 sessions.

### Behavioral analysis

We analyzed reaches from the behavioral trials. All successful trials from all sessions were labeled with reach direction (right, down, left, or up) and stimulation condition (stimulation or no stimulation). Data groups were compared statistically by Mann-Whitney U-test of the distribution of reach times. Our experimental design focused on examining how stimulation would affect behavioral metrics, including reach time, path length and direction, which we carefully recorded. For these hypotheses, we did not adjust p-values as the nature of our analysis were not exploratory.

We tracked the reach path and reach time and analyzed movement in three-dimensional space for both stimulation and control (no stimulation) conditions. We measured the time between the go-tone and the finger’s entry into the end target, and we call this time the “reach time.”

### Behavioral data preprocessing

All reach trials were aggregated over all dates and conditions for each monkeys individually. Trials were excluded from analysis if the reach was unsuccessful or if the trail was a warmup trial as judged from a wait time before the go-tone <0.3 s, or a circle size >50 pixels radius (approx. 25 mm diameter). The counts of included trials are listed in Supp. Table 1.

The neural data were screened to remove malfunctioning electrodes by measuring the impedance of the electrodes and by visual inspection of the voltage traces. Additionally, the standard deviation of the raw neural signal was computed for each trial, and if the z-score of the standard deviation was >6, the trial was excluded.

### Neural data analysis

Neural activity was recorded during behavior and stimulation (1 kHz sampling frequency, Grapevine Nomad, Ripple Neuro). All neural data analyses were conducted separately for the two monkeys. High impedance electrodes were excluded from analyses.

The process involved segregating all trials into stimulation and non-stimulation trials. Each trial was then split into three epochs: before stimulation, during stimulation (or during when the stimulation would have occurred), and after stimulation. We analyzed 500 ms of local field potential (LFP) activity immediately preceding the onset of stimulation (“before stimulation”). The “during stimulation” period encompassed the 900 ms of LFP recorded during stimulation, and the “after stimulation” period covered the 900 ms following the end of stimulation. Artifact removal was applied to each epoch (see “Photo-Induced Artifact Removal” section below).

After removing photo-induced artifact, we quantified the effect of stimulation by analyzing the time-frequency content of the data. Multitaper spectrograms of ECoG data from each trial were calculated with 50-ms non-overlapping windows, five tapers and a 128-sample length Fast Fourier Transform (FFT) between 1-200 Hz. The values of each bin are normalized by the 500-ms pre-stimulation activity. The 50 ms immediately following the start and termination of stimulation are excluded from analysis to avoid including any residual artifact in the signal at these times.

To quantify the changes due to stimulation, we calculated the average values of the spectrogram per frequency bin. Spectrograms were normalized to the 500 ms preceding stimulation and the resulting values were used to determine if statistically significant changes were induced before, during, and after stimulation using the Mann-Whitney U test. For control experiments, the data processing matched the stimulated condition, although the stimulation did not occur.

### Photo-induced artifact removal

To isolate photo-induced artifacts from the electrocorticography acquired during simultaneous optical stimulation in vivo, we generated a dataset of these artifacts, absent any neural signals, by optically stimulating an electrode array submerged in a saline bath. We modeled the photo-induced artifact using these saline data, fit the model to the in vivo ECoG, and removed the estimated photo-induced artifact. The procedure involved the following steps (Supp. Fig. 8):

1. The time surrounding each stimulation in the saline data was isolated, from 100 ms before the onset of stimulation to one second after the end of stimulation. Each of the 32 channels were stacked with their 100 corresponding stimulation pulses to form a matrix (32 × 100 × length of time).
2. The matrix was split along the time axis into three matrices corresponding to before stimulation, during stimulation, and after stimulation.
3. The saline data at the time of stimulation onset and offset (data point time = 0 for matrices “during stimulation” and “after stimulation,” respectively) were set to 0 mV and the whole traces were shifted accordingly.
4. Photo-induced artifact was modeled from these data using principal component analysis (PCA) conducted separately on the “during stimulation” and “after stimulation” matrices, with the top three components extracted from each.

This model was subsequently employed to eliminate the artifact from the in vivo data as follows:

5. The in vivo data was isolated and stacked in the same way as the saline data, such that a data matrix (channels × pulses × length of time), with identical time length, was formed.
6. The matrix was split along the time axis into three matrices corresponding to before stimulation during stimulation, and after stimulation.
7. The ECoG data at the time of stimulation onset and offset (data point time = 0 for matrices “during stimulation” and “after stimulation”) were set to 0 mV and the whole traces were shifted accordingly.
8. The ECoG data from the “during stimulation” and “after stimulation” matrices were projected onto the corresponding models derived from saline, and the estimated photo-induced artifact was regressed out of the NHP data using linear regression.
9. The three matrices were rejoined along the time axis, adding an offset to the later matrices such that the average LFP value of the last 50 ms of the preceding matrix and the first 50 ms of the later matrix had the same average value.

This procedure enabled the separation of the photo-induced artifact from neural activity, and retaining the neural activity in the recorded signal (Fig. 4; Supp. Fig. 8), as confirmed by controls (Fig. 4).

### Simulation

We simulated large-scale neural spiking activity and LFPs across cortical layers using the extended VERTEX model, which allows for optogenetic stimulation and neural activity recording ^56^. The tissue model was 1.51×11.51×12.61mm deep with LFP recording sites arranged in a 31×131×16 grid, and a 200 µm optical fiber placed orthogonally on the center of the cortical surface (Supp. Fig. 9). We used the VERTEX Jaws opsin model, which hyperpolarizes neuronal membrane currents when optically evoked. Optogenetic inhibition was modeled with a train of optical pulses matching our experimental conditions (594 nm wavelength, 10 pulses, 9001ms on/29001ms off). To examine local firing rates around the optic fiber, Supp. Fig. 5a, b show neuron firing rates along the z-axis within a 400 µm cylinder around the optical fiber. Supp. Fig. 5c shows the time-frequency spectrogram of the LFP recording sites centered in each layer.

## References

1. Camporeze, B. et al. Optogenetics: the new molecular approach to control functions of neural cells in epilepsy, depression and tumors of the central nervous system. Am J Cancer Res 8, 1900–1918 (2018).

2. Carter, M. E. & de Lecea, L. Optogenetic investigation of neural circuits in vivo. Trends Mol Med 17, 197–206 (2011).

3. Barnett, S. C., Perry, B. A. L., Dalrymple-Alford, J. C. & Parr-Brownlie, L. C. Optogenetic stimulation: Understanding memory and treating deficits. Hippocampus 28, 457–470 (2018).

4. Adhikari, A. et al. Basomedial amygdala mediates top-down control of anxiety and fear. Nature 527, 179–185 (2015).

5. Chen, B. T. et al. Rescuing cocaine-induced prefrontal cortex hypoactivity prevents compulsive cocaine seeking. Nature 496, 359–362 (2013).

6. Yang, X. et al. A novel mechanism of memory loss in Alzheimer’s disease mice via the degeneration of entorhinal-CA1 synapses. Mol Psychiatry 23, 199–210 (2018).

7. Jiang, W., Tremblay, F. & Elaine Chapman, C. Context-dependent tactile texture-sensitivity in monkey M1 and S1 cortex. J Neurophysiol 120, 2334–2350 (2018).

8. Gradinaru, V., Mogri, M., Thompson, K. R., Henderson, J. M. & Deisseroth, K. Optical deconstruction of parkinsonian neural circuitry. Science (1979) 324, 354–359 (2009).

9. Yazdan-Shahmorad, A., Silversmith, D. B., Kharazia, V. & Sabes, P. N. Targeted cortical reorganization using optogenetics in non-human primates. Elife 7, 1–21 (2018).

10. Jagust, W. Vulnerable Neural Systems and the Borderland of Brain Aging and Neurodegeneration. Neuron 77, 219–234 (2013).

11. Uhlhaas, P. J. & Singer, W. Neural Synchrony in Brain Disorders: Relevance for Cognitive Dysfunctions and Pathophysiology. Neuron 52, 155–168 (2006).

12. Tønnesen, J. Optogenetic cell control in experimental models of neurological disorders. Behavioural Brain Research 255, 35–43 (2013).

13. Ruggiero, R. N. et al. Cannabinoids and vanilloids in schizophrenia: Neurophysiological evidence and directions for basic research. Front Pharmacol 8, 1–27 (2017).

14. Roelfsema, P. R. & Treue, S. Basic neuroscience research with nonhuman primates: A small but indispensable component of biomedical research. Neuron Preprint at 10.1016/j.neuron.2014.06.003 (2014).

15. Lear, A. et al. Understanding them to understand ourselves: The importance of NHP research for translational neuroscience. Current Research in Neurobiology 100049 (2022).

16. Diester, I. et al. An optogenetic toolbox designed for primates. Nat Neurosci 14, 387–397 (2011).

17. Han, X. et al. Millisecond-Timescale Optical Control of Neural Dynamics in the Nonhuman Primate Brain. Neuron 62, 191–198 (2009).

18. Renz, A. F., et al. Opto-E-Dura: A Soft, Stretchable ECoG Array for Multimodal, Multiscale Neuroscience. Adv Healthc Mater 9, 1–11 (2020).

19. Richner, T. J. et al. Optogenetic micro-electrocorticography for modulating and localizing cerebral cortex activity. J Neural Eng 11, (2014).

20. Komatsu, M., Sugano, E., Tomita, H. & Fujii, N. A chronically implantable bidirectional neural interface for non-human primates. Front Neurosci 11, 1–9 (2017).

21. Macknik, S. L. et al. Advanced Circuit and Cellular Imaging Methods in Nonhuman Primates. J Neurosci 39, 8267–8274 (2019).

22. Slovin, H., Arieli, A., Hildesheim, R. & Grinvald, A. Long-term voltage-sensitive dye imaging reveals cortical dynamics in behaving monkeys. J Neurophysiol 88, 3421–3438 (2002).

23. Arieli, A., Grinvald, A. & Slovin, H. Dural substitute for long-term imaging of cortical activity in behaving monkeys and its clinical implications. J Neurosci Methods 114, 119–133 (2002).

24. Ruiz, O. et al. Optogenetics through windows on the brain in the nonhuman primate. J Neurophysiol 110, 1455–1467 (2013).

25. Yazdan-Shahmorad, A. et al. A Large-Scale Interface for Optogenetic Stimulation and Recording in Nonhuman Primates. Neuron 89, 927–939 (2016).

26. Chernov, M. & Roe, A. W. Infrared neural stimulation: a new stimulation tool for central nervous system applications. Neurophotonics 1, 011011 (2014).

27. Zaraza, D., et al. Going wireless: an optical imaging and optogenetics system for use in awake behaving primates. 1122705, (2020).

28. Griggs, D. J., Belloir, T. & Yazdan-Shahmorad, A. Large-scale neural interfaces for optogenetic actuators and sensors in non-human primates. SPIE BiOS 1166305, 17 (2021).

29. Yazdan-Shahmorad, A. et al. Demonstration of a setup for chronic optogenetic stimulation and recording across cortical areas in non-human primates. SPIE BiOS 9305, 93052K (2015).

30. Krauze, M. T. et al. Reflux-free cannula for convection-enhanced high-speed delivery of therapeutic agents. J Neurosurg 103, 923–929 (2005).

31. Yazdan-Shahmorad, A. et al. A Large-Scale Interface for Optogenetic Stimulation and Recording in Nonhuman Primates. Neuron 89, 927–939 (2016).

32. Khateeb, K., Griggs, D. J., Sabes, P. N. & Yazdan-Shahmorad, A. Convection Enhanced Delivery of Optogenetic Adeno-associated Viral Vector to the Cortex of Rhesus Macaque Under Guidance of Online MRI Images. JoVE e59232 (2019) doi:doi:10.3791/59232.

33. Tremblay, S. et al. An Open Resource for Non-human Primate Optogenetics. Neuron 1–16 (2020) doi:10.1016/j.neuron.2020.09.027.

34. Rajalingham, R. et al. Chronically implantable LED arrays for behavioral optogenetics in primates. Nat Methods 18, 1112–1116 (2021).

35. Pollmann, E. H. et al. A subdural CMOS optical device for bidirectional neural interfacing. Nat Electron 7, 829–841 (2024).

36. Khanna, P. et al. Low-frequency stimulation enhances ensemble co-firing and dexterity after stroke. Cell 184, 912–930.e20 (2021).

37. Griggs, D. J. Accessible Optogenetic Technologies for Non-Human Primate Research. ProQuest Dissertations and Theses (University of Washington PP - United States -- Washington, United States -- Washington, 2022).

38. Bloch, J. Computational and Experimental Advances in Clinically Relevant Neurostimulation of Non-Human Primates. ProQuest Dissertations and Theses (University of Washington PP - United States -- Washington, United States -- Washington, 2023).

39. Ojemann, W. K. S. et al. A mri-based toolbox for neurosurgical planning in nonhuman primates. Journal of Visualized Experiments 2020, 1–16 (2020).

40. Iritani, R. et al. A Neural Implant Design Toolbox for Nonhuman Primates. Journal of Visualized Experiments (2024) doi:10.3791/66167.

41. Griggs, D. J. et al. Multi-modal artificial dura for simultaneous large-scale optical access and large-scale electrophysiology in non-human primate cortex. Journal of Neural Engineering, Special Issue ‘Neuroelectronic Interfaces’ 18, 2021.02.03.429596 (2021).

42. Griggs, D. J., et al. Optimized large-scale optogenetic interface for non-human primates. SPIE BiOS 1086605, 3 (2019).

43. Griggs, D. J. et al. Demonstration of an Optimized Large-scale Optogenetic Cortical Interface for Non-human Primates. Ieee 119395, 1–4 (2022).

44. Ozden, I. et al. A coaxial optrode as multifunction write-read probe for optogenetic studies in non-human primates. J Neurosci Methods 219, 142–154 (2013).

45. McAlinden, N. et al. Thermal and optical characterization of micro-LED probes for in vivo optogenetic neural stimulation. Opt Lett 38, 992–994 (2013).

46. Andersen, P. & Moser, E. I. Brain temperature and hippocampal function. Hippocampus 5, 491–498 (1995).

47. Price, R. R. The AAPM/RSNA physics tutorial for residents: MR imaging safety considerations. Radiographics 19, 1641–1651 (1999).

48. Galvan, A. et al. Nonhuman Primate Optogenetics: Recent Advances and Future Directions. The Journal of Neuroscience 37, 10894 (2017).

49. Acker, L., Pino, E. N., Boyden, E. S. & Desimone, R. FEF inactivation with improved optogenetic methods. Proceedings of the National Academy of Sciences 113, E7297–E7306 (2016).

50. Khateeb, K., Griggs, D. J., Sabes, P. N. & Yazdan-Shahmorad, A. Convection Enhanced Delivery of Optogenetic Adeno-associated Viral Vector to the Cortex of Rhesus Macaque Under Guidance of Online MRI Images. J Vis Exp 1–8 (2019) doi:10.3791/59232.

51. Griggs, D. J. et al. Improving the efficacy and accessibility of intracranial viral vector delivery in non-human primates. Pharmaceutics 2022.06.06.494543 (2022) doi:10.1101/2022.06.06.494543.

52. Chuong, A. S. et al. Noninvasive optical inhibition with a red-shifted microbial rhodopsin. Nat Neurosci 17, 1123–1129 (2014).

53. Acker, L., Pino, E. N., Boyden, E. S. & Desimone, R. FEF inactivation with improved optogenetic methods. Proc Natl Acad Sci U S A 113, E7297–E7306 (2016).

54. Tomsett, R. J. et al. Virtual Electrode Recording Tool for EXtracellular potentials (VERTEX): comparing multi-electrode recordings from simulated and biological mammalian cortical tissue. Brain Struct Funct 220, 2333–2353 (2015).

55. Binzegger, T., Douglas, R. J. & Martin, K. A. C. A quantitative map of the circuit of cat primary visual cortex. Journal of Neuroscience 24, 8441–8453 (2004).

56. Pierce, A. F., Shupe, L., Bloch, J., Fetz, E. & Yazdan-Shahmorad, A. Flexible modeling of large-scale neural network stimulation: Electrical and optical extensions to The Virtual Electrode Recording Tool for EXtracellular Potentials (VERTEX). J Neurosci Methods 422, 110514 (2025).

57. Stanis, N., Griggs, D., Bloch, J. & Yazdan-Shahmorad, A. Neural and Behavioral Perturbations via Optogenetic Inhibition of the Posterior Parietal Cortex during Motor Planning. in EMBC 2025: 47th Annual International Conference of the IEEE Engineering in Medicine and Biology Society (accepted for publication) (Copenhagen, Denmark, 2025).

58. Khateeb, K. et al. A versatile toolbox for studying cortical physiology in primates. Cell Reports Methods 100183 (2022) doi:10.1016/j.crmeth.2022.100183.

59. Zhou, J. et al. Neuroprotective Effects of Electrical Stimulation Following Ischemic Stroke in Non-Human Primates *. IEEE EMBC 119395, 1–4 (2022).

60. Schwock, F. et al. Inferring Neural Communication Dynamics from Field Potentials Using Graph Diffusion Autoregression. bioRxiv 2024.02.26.582177 (2024) doi:10.1101/2024.02.26.582177.

61. Zhou, J., Khateeb, K. & Yazdan-Shahmorad, A. Early Intervention with Electrical Stimulation Reduces Neural Damage After Stroke in Non-human Primates. bioRxiv 2023.12.18.572235 (2023) doi:10.1101/2023.12.18.572235. (now in press with Nature Communications)

62. Ohayon, S., Grimaldi, P., Schweers, N. & Tsao, D. Y. Saccade modulation by optical and electrical stimulation in the macaque frontal eye field. Journal of Neuroscience 33, 16684–16697 (2013).

63. Hart, W. L. et al. Combined optogenetic and electrical stimulation of auditory neurons increases effective stimulation frequency - An in vitro study. J Neural Eng 17, (2020).

64. Vargo, S. M. et al. Smart Dura: A Monolithic Optoelectrical Surface Array for Neural Interfacing with Primate Cortex. in 2023 11th International IEEE/EMBS Conference on Neural Engineering (NER) 1–6 (2023). doi:10.1109/NER52421.2023.10123755.

65. Belloir, T. et al. Large-scale multimodal surface neural interfaces for primates. iScience 26, 105866 (2023).

66. Khodagholy, D. et al. NeuroGrid: recording action potentials from the surface of the brain. Nat Neurosci 18, 310–315 (2015).

67. Montalvo Vargo, S., et al. Smart Dura: a functional artificial dura for multimodal neural recording and modulation. bioRxiv 2025.02.26.640369 (2025) doi:10.1101/2025.02.26.640369.

68. Ji, B. et al. Flexible polyimide-based hybrid opto-electric neural interface with 16 channels of micro-LEDs and electrodes. Microsyst Nanoeng 4, (2018).

69. Reddy, J. W., Kimukin, I., Towe, E. & Chamanzar, M. Flexible, Monolithic, High-Density μlED Neural Probes for Simultaneous Optogenetics Stimulation and Recording. International IEEE/EMBS Conference on Neural Engineering, NER 2019-March, 831–834 (2019).

70. Clark, A. M. et al. An optrode array for spatiotemporally-precise large-scale optogenetic stimulation of deep cortical layers in non-human primates. Commun Biol 7, (2024).

71. Gong, X. et al. An Ultra-Sensitive Step-Function Opsin for Minimally Invasive Optogenetic Stimulation in Mice and Macaques. Neuron 107, 38–51.e8 (2020).

72. Chen, L. M. et al. A chamber and artificial dura method for long-term optical imaging in the monkey. J Neurosci Methods 113, 41–49 (2002).

73. Chiang, C.-H. et al. Development of a neural interface for high-definition, long-term recording in rodents and nonhuman primates. Sci Transl Med 12, eaay4682 (2020).

74. Yao, Z. & Yazdan-Shahmorad, A. A Quantitative Model for Estimating the Scale of Photochemically Induced Ischemic Stroke. Conf Proc IEEE Eng Med Biol Soc 2018, 2744–2747 (2018).

75. Khateeb, K. et al. A Practical Method for Creating Targeted Focal Ischemic Stroke in the Cortex of Nonhuman Primates(.). Conf Proc IEEE Eng Med Biol Soc 2019, 3515–3518 (2019).

76. Eldridge, M. A. G. & Galvan, A. Vectorology for Optogenetics and Chemo-Genetics. http://www.springer.com/series/7657 (2023).

77. Griggs, D. J. et al. Improving the Efficacy and Accessibility of Intracranial Viral Vector Delivery in Non-Human Primates. Pharmaceutics 14, (2022).

78. Meyer, H. S. et al. Inhibitory interneurons in a cortical column form hot zones of inhibition in layers 2 and 5A. Proc Natl Acad Sci U S A 108, 16807–16812 (2011).

79. Kooijmans, R. N., Sierhuis, W., Self, M. W. & Roelfsema, P. R. A Quantitative Comparison of Inhibitory Interneuron Size and Distribution between Mouse and Macaque V1, Using Calcium-Binding Proteins. Cereb Cortex Commun 1, 1–14 (2020).

80. Thio, B. J. & Grill, W. M. Relative contributions of different neural sources to the EEG. Neuroimage 275, 120179 (2023).

81. Baratham, V. L. et al. Columnar Localization and Laminar Origin of Cortical Surface Electrical Potentials. Journal of Neuroscience 42, 3733–3748 (2022).

82. Yazdan-Shahmorad, A., Kipke, D. R. & Lehmkuhle, M. J. Polarity of cortical electrical stimulation differentially affects neuronal activity of deep and superficial layers of rat motor cortex. Brain Stimul 4, 228–241 (2011).

83. Yazdan-Shahmorad, A., Kipke, D. R. & Lehmkuhle, M. J. High gamma power in ECoG reflects cortical electrical stimulation effects on unit activity in layers V/VI. J Neural Eng 10, 066002 (2013).

84. Yazdan-Shahmorad, A., Kipke, D. R. & Lehmkuhle, M. J. Polarity of cortical electrical stimulation differentially affects neuronal activity of deep and superficial layers of rat motor cortex. Brain Stimul 4, 228–241 (2011).

85. Jacques, S. L. Erratum: Optical properties of biological tissues: A review (Physics in Medicine and Biology (2013) 58). Phys Med Biol 58, 5007–5008 (2013).

86. Watanabe, H. et al. Forelimb movements evoked by optogenetic stimulation of the macaque motor cortex. Nat Commun 11, 1–9 (2020).

87. O’Shea, D. J. et al. Direct neural perturbations reveal a dynamical mechanism for robust computation. bioRxiv 2022.12.16.520768 (2022) doi:10.1101/2022.12.16.520768.

88. Fetsch, C. R. et al. Focal optogenetic suppression in macaque area MT biases direction discrimination and decision confidence, but only transiently. Elife 7, 1–23 (2018).

89. Katz, L. N. et al. Optogenetic Manipulation of Covert Attention in the Nonhuman Primate. J Cogn Neurosci 1–20 (2024) doi:10.1162/jocn_a_02274.

90. Mountcastle, V. B., Lynch, J. C., Georgopoulos, A., Sakata, H. & Acuna, C. Posterior parietal association cortex of the monkey: command functions for operations within extrapersonal space. J Neurophysiol 38, 871–908 (1975).

91. Kalaska, J. F., Cohen, D. A., Prud’homme, M. & Hyde, M. L. Parietal area 5 neuronal activity encodes movement kinematics, not movement dynamics. Exp Brain Res 80, 351–364 (1990).

92. Kalaska, J. F. & Crammond, D. J. Deciding not to GO: neuronal correlates of response selection in a GO/NOGO task in primate premotor and parietal cortex. Cereb Cortex 5, 410–428 (1995).

93. Lacquaniti, F., Guigon, E., Bianchi, L., Ferraina, S. & Caminiti, R. Representing spatial information for limb movement: role of area 5 in the monkey. Cereb Cortex 5, 391–409 (1995).

94. Mountcastle, V. B. The parietal system and some higher brain functions. Cereb Cortex 5, 377–390 (1995).

95. Caminiti, R., Ferraina, S. & Johnson, P. B. The sources of visual information to the primate frontal lobe: a novel role for the superior parietal lobule. Cereb Cortex 6, 319–328 (1996).

96. Johnson, P. B., Ferraina, S., Bianchi, L. & Caminiti, R. Cortical networks for visual reaching: physiological and anatomical organization of frontal and parietal lobe arm regions. Cereb Cortex 6, 102–119 (1996).

97. Battaglia-Mayer, A. et al. Early coding of reaching in the parietooccipital cortex. J Neurophysiol 83, 2374–2391 (2000).

98. Battaglia-Mayer, A., Caminiti, R., Lacquaniti, F. & Zago, M. Multiple levels of representation of reaching in the parieto-frontal network. Cereb Cortex 13, 1009–1022 (2003).

99. Andersen, R. A. & Buneo, C. A. Intentional maps in posterior parietal cortex. Annu Rev Neurosci 25, 189–220 (2002).

100. Buneo, C. A., Jarvis, M. R., Batista, A. P. & Andersen, R. A. Direct visuomotor transformations for reaching. Nature 416, 632–636 (2002).

101. Cohen, Y. E. & Andersen, R. A. A common reference frame for movement plans in the posterior parietal cortex. Nat Rev Neurosci 3, 553–562 (2002).

102. Koch, G. et al. Functional interplay between posterior parietal and ipsilateral motor cortex revealed by twin-coil transcranial magnetic stimulation during reach planning toward contralateral space. Journal of Neuroscience 28, 5944–5953 (2008).

103. Snyder, L. H., Batista, A. P. & Andersen, R. A. Coding of intention in the posterior parietal cortex. Nature 386, 167–170 (1997).

104. Tanné-Gariépy, J., Rouiller, E. M. & Boussaoud, D. Parietal inputs to dorsal versus ventral premotor areas in the macaque monkey: evidence for largely segregated visuomotor pathways. Exp Brain Res 145, 91–103 (2002).

105. Croxson, P. L. et al. Quantitative investigation of connections of the prefrontal cortex in the human and macaque using probabilistic diffusion tractography. J Neurosci 25, 8854–8866 (2005).

106. Makris, N. et al. Segmentation of subcomponents within the superior longitudinal fascicle in humans: a quantitative, in vivo, DT-MRI study. Cereb Cortex 15, 854–869 (2005).

107. Rushworth, M. F. S. & Taylor, P. C. J. TMS in the parietal cortex: updating representations for attention and action. Neuropsychologia 44, 2700–2716 (2006).

108. Rozzi, S. et al. Cortical connections of the inferior parietal cortical convexity of the macaque monkey. Cereb Cortex 16, 1389–1417 (2006).

109. Wise, S. P., Boussaoud, D., Johnson, P. B. & Caminiti, R. Premotor and parietal cortex: Corticocortical connectivity and combinatorial computations. Annu Rev Neurosci 20, 25–42 (1997).

110. Goldring, A. B., Cooke, D. F., Pineda, C. R., Recanzone, G. H. & Krubitzer, L. A. Functional characterization of the fronto-parietal reaching and grasping network: reversible deactivation of M1 and areas 2, 5, and 7b in awake behaving monkeys. J Neurophysiol 127, 1363–1387 (2022).

111. Li, Y., Wang, Y. & Cui, H. Posterior parietal cortex predicts upcoming movement in dynamic sensorimotor control. Proceedings of the National Academy of Sciences 119, e2118903119 (2022).

112. Churchland, M. M. & Shenoy, K. V. Delay of movement caused by disruption of cortical preparatory activity. J Neurophysiol 97, 348–359 (2007).

113. Busan, P. et al. Effect of transcranial magnetic stimulation (TMS) on parietal and premotor cortex during planning of reaching movements. PLoS One 4, (2009).

114. Konen, C. S., Mruczek, R. E. B., Montoya, J. L. & Kastner, S. Functional organization of human posterior parietal cortex: grasping-and reaching-related activations relative to topographically organized cortex. J Neurophysiol 109, 2897–2908 (2013).

115. Bloch, J., Shea-brown, E., Harchaoui, Z. & Shojaie, A. iScience ll Network structure mediates functional reorganization induced by optogenetic stimulation of non-human primate sensorimotor cortex. iScience 25, 104285 (2022).

116. Bloch, J. A. et al. Cortical Stimulation Induces Network-Wide Coherence Change in Non-Human Primate Somatosensory Cortex∗. Proceedings of the Annual International Conference of the IEEE Engineering in Medicine and Biology Society, EMBS 6446–6449 (2019) doi:10.1109/EMBC.2019.8856633.

117. Zhou, A. et al. A wireless and artefact-free 128-channel neuromodulation device for closed-loop stimulation and recording in non-human primates. Nat Biomed Eng 3, 15–26 (2019).

118. Belloir, T., et al. iScience Large-scale multimodal surface neural interfaces for non-human primates. iScience (2022).

119. Vargo, S. M., Belloir, T., Kimukin, I., Ahmed, Z. & Griggs, D. Smart Dura1: A monolithic optoelectrical surface array for neural interfacing with primate cortex. EMBC NER 116464, 2–6 (2023).

120. Qiao, N., Ma, L., Zhang, Y. & Wang, L. Update on Nonhuman Primate Models of Brain Disease and Related Research Tools. Biomedicines 11, 2516 (2023).

121. Brainard, D. H. The Psychophysics Toolbox. Spat Vis 10, 433–436 (1997).

122. Pelli, D. G. The VideoToolbox software for visual psychophysics: Transforming numbers into movies. Spat Vis 10, 437–442 (1997).

123. Kleiner, M., Brainard, D. H. & Pelli, D. G. What’s new in Psychtoolbox-3? in Perception 36 ECVP Abstract Supplement (2007).

124. Griggs, D. J. et al. Autonomous cage-side system for remote training of non-human primates. J Neurosci Methods 348, 108969 (2020).

