## Supplemental Data for "A large-scale optogenetic neurophysiology platform for improving accessibility in non-human primate behavioral experiments"

### Supplemental Figures

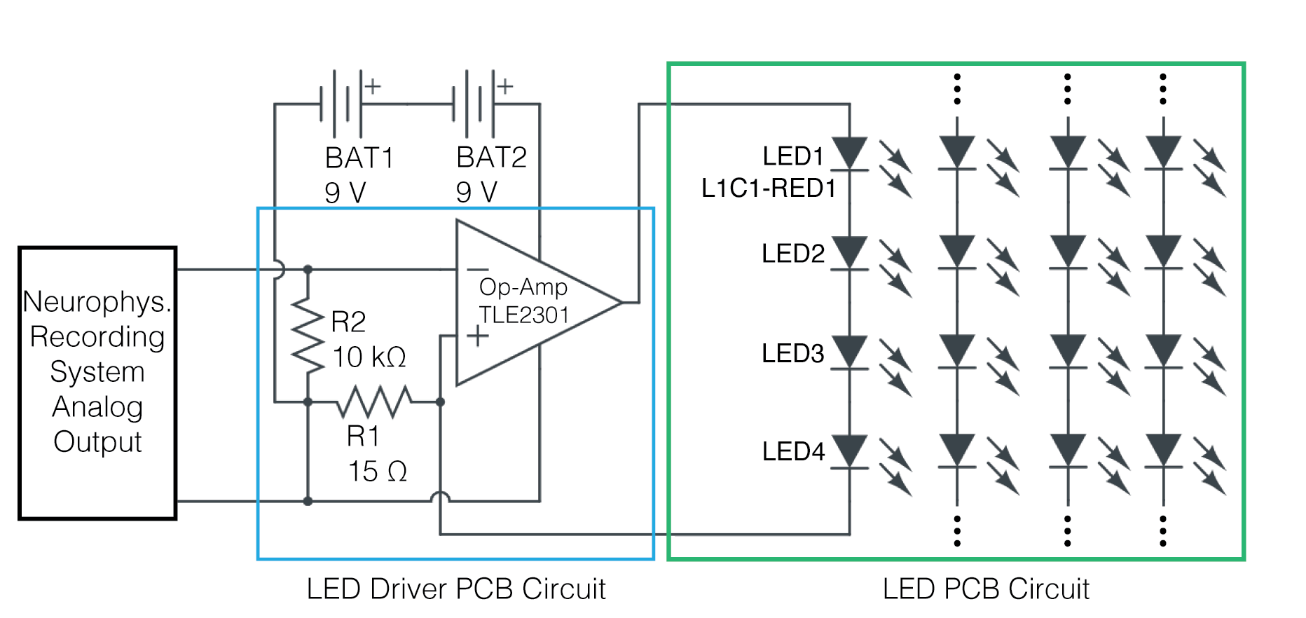

Supp. Fig. 1: Diagram of LED driving circuitry for the 4 × 4 array, which has four individually drivable rows of four LEDs each. The 3 × 5 array, which has 15 individually drivable LEDs, has identical circuitry except each LED Driver PCB Circuit drives only one LED, rather than four in series. Figure adapted from ^39^ with permission from IEEE EMBC.

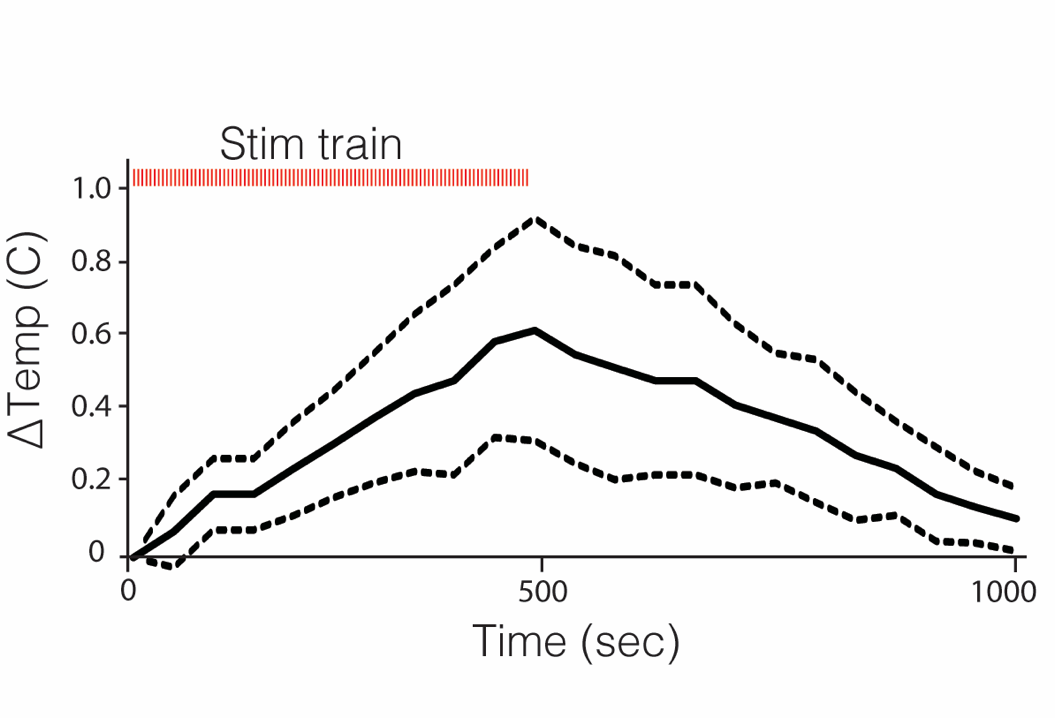

*Supp. Fig. 2: Temperature change from optical stimulation. Stimulation protocol: 100 pulses of 900ms pulse width, 5 second period. Mean and standard deviation of temperature change at each time point during and after stimulation protocol is shown. Temperature started at room temp and each measurement was recorded at pulse offset. Experimental stack up includes: MMAD, stimulation holder, cover glass, tube, and the 4x4 LED array (12 mW/mm^2^) on top of a temperature probe in agarose hydrogel to mimic the brain.*

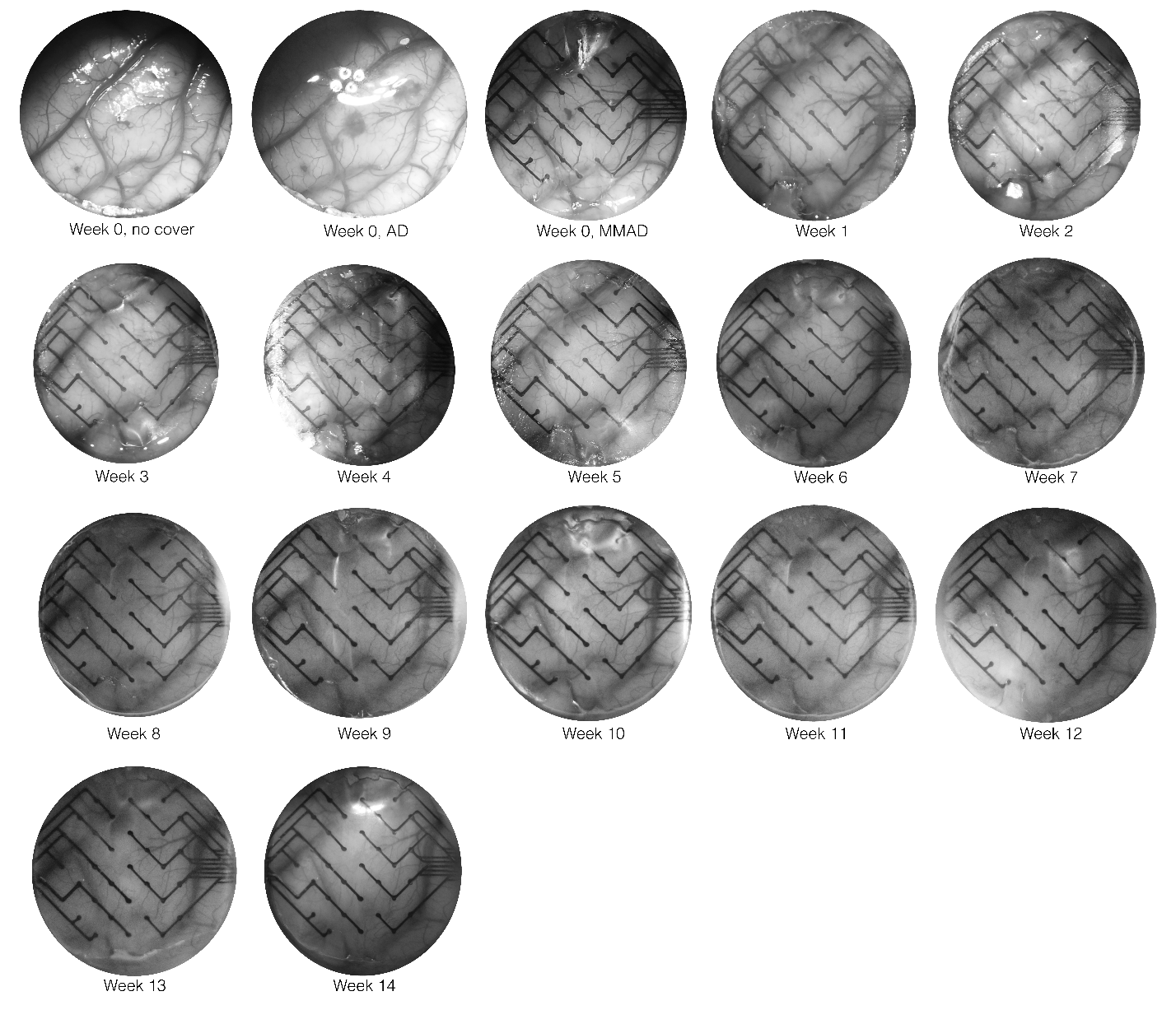

*Supp. Fig. 3: Optical access over 14 weeks in Monkey H.*

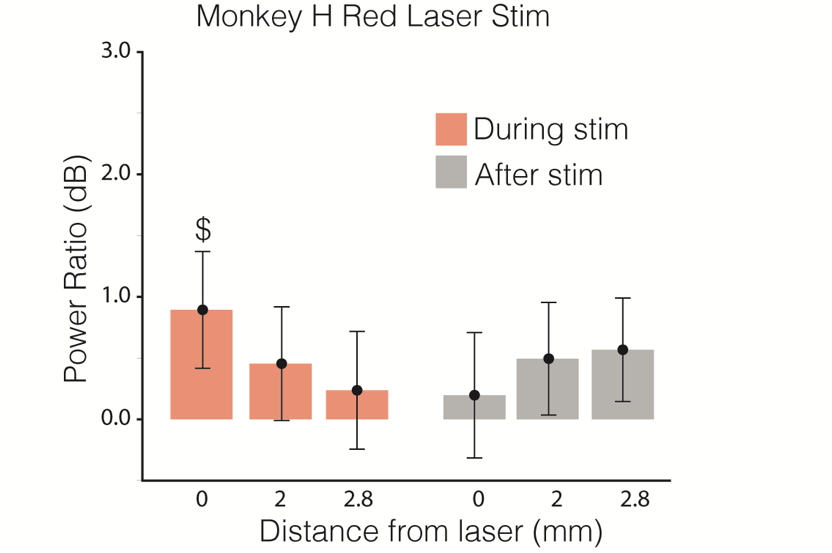

*Supp. Fig. 4: Laser stimulation during rest in Monkey H. The power ratios of responses are calculated by taking the ECoG power during stimulation and dividing by the ECoG power from before stimulation (1-200Hz), then converting to decibels. Whiskers are 95% confidence intervals of the medians generated with a bootstrapping method, and asterisks indicate significant differences from zero (p < 0.05). Single red laser stimulation was performed with three conditions: illumination directly over, 2.0 mm from, and 2.8 mm from the recording channel.*

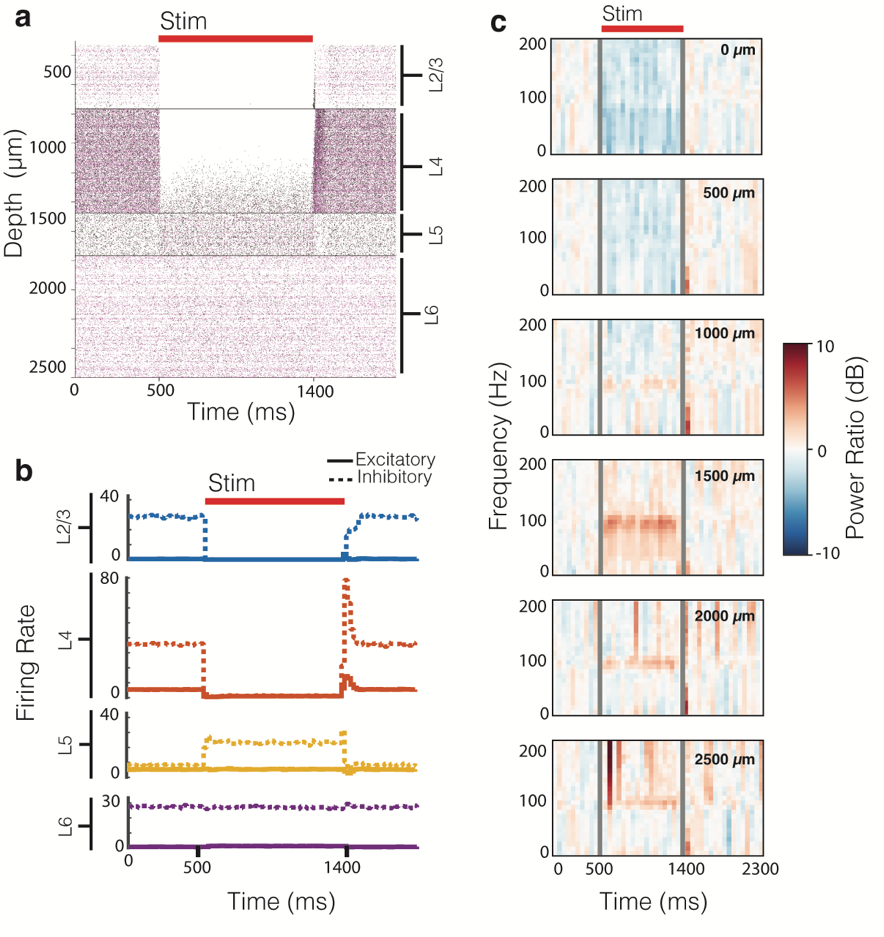

*Supp. Fig. 5: Simulation of effects surface optogenetic inhibition on cortical layers.* ***(a)*** *Raster of spiking excitatory (black ticks) and inhibitory (magenta ticks) neurons at different cortical depths before, during, and after stimulation.* ***(b)*** *Firing rate of excitatory (solid line) and inhibitory (dashed line) neurons of (a), categorized by cortical layer.* ***(c)*** *Spectrogram of LFPs at different depths of the cortical tissue. Grey bars indicate bins during the first 50 ms of stimulation onset and offset – data during these times are excluded in replica of Fig. 5 results. Inhibition is seen in upper layers of cortex during stimulation while lower layers show increased activity.*

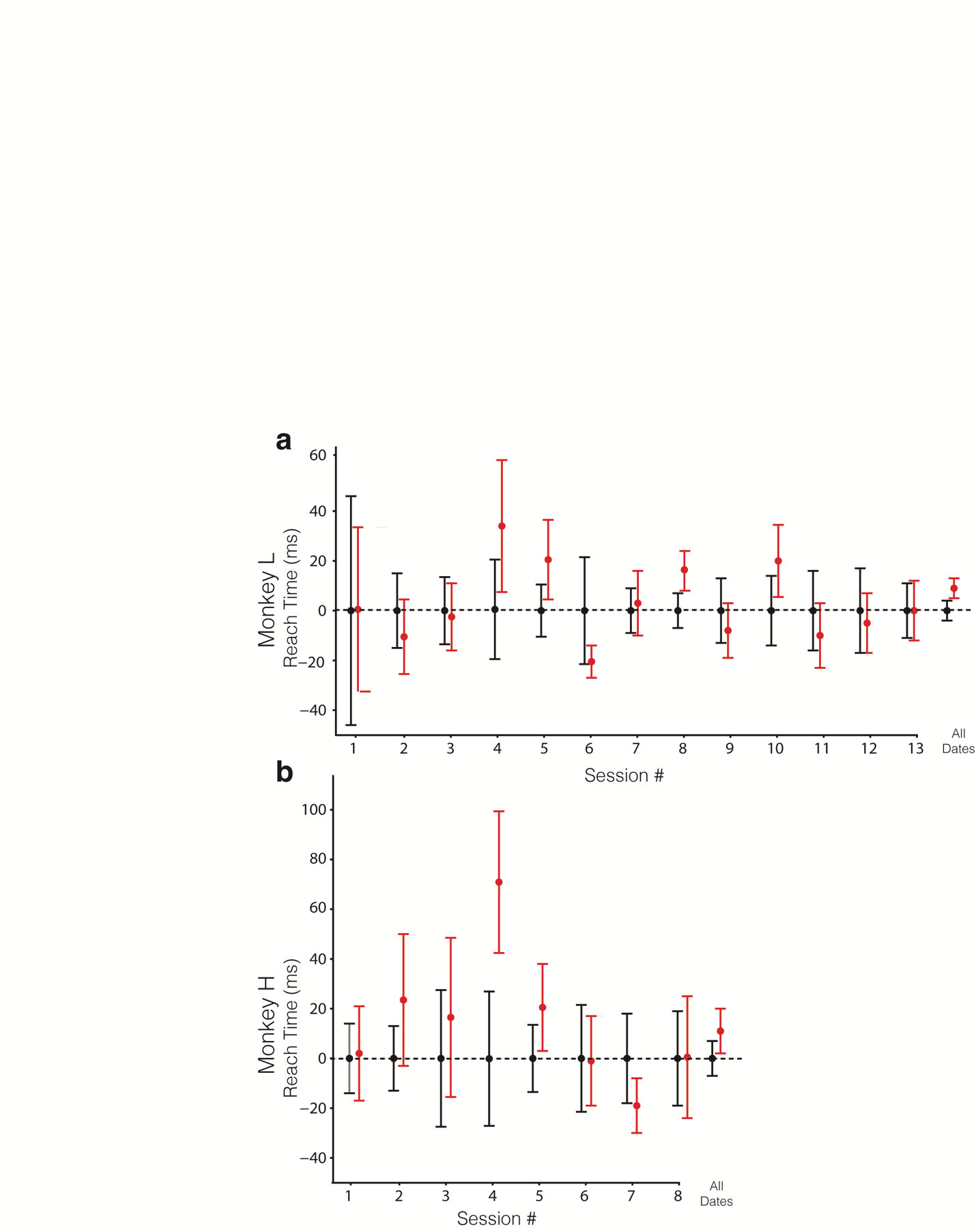

*Supp. Fig. 6: Difference in reach time for (a) Monkey L and (b) Monkey H. Red shows stim trials and black shows control trials. Reach time of individual sessions is normalized to the non-stimulated condition. Median and 95% confidence intervals are reported.*

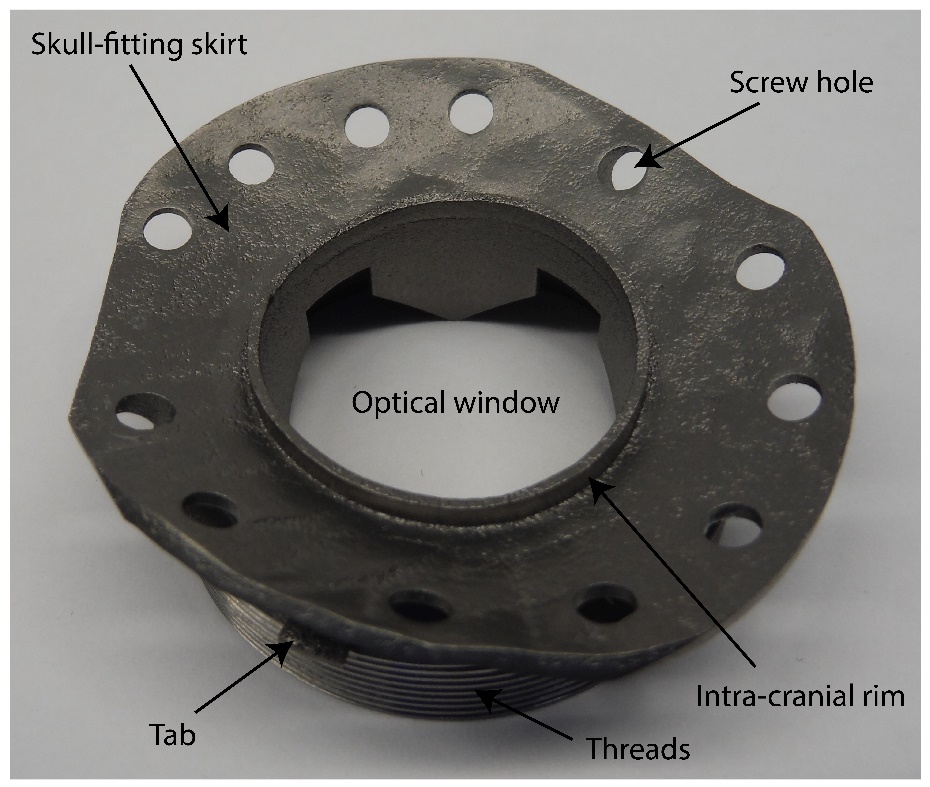

*Supp. Fig. 7: Chamber viewed from below skirt. Figure adapted from ^40^ with permission from IEEE EMBC.*

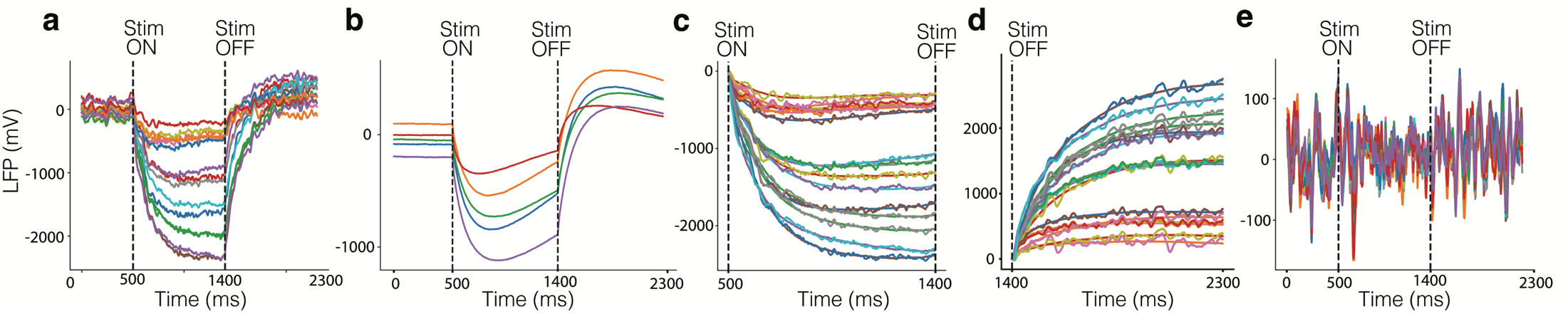

*Supp. Fig.8: Method of light-induced artifact removal.* ***(a)*** *Example single trial LFP timeseries of different electrodes during optical stimulation of cortex. Electrodes were over both opsin expressing and opsin deficient areas.* *High impedance electrodes were excluded.* ***(b)*** *Example recording during optical stimulation of ECoG array in a saline bath during a single trial.* ***(c, d)*** *Neural recordings overlaid with a reconstructed artifact template found by projecting the recording into the first four principal components of saline recording during (c) and after (d) stimulation.* ***(e)*** *LFP timeseries with photo-artifact removed using the reconstructed artifact template.*

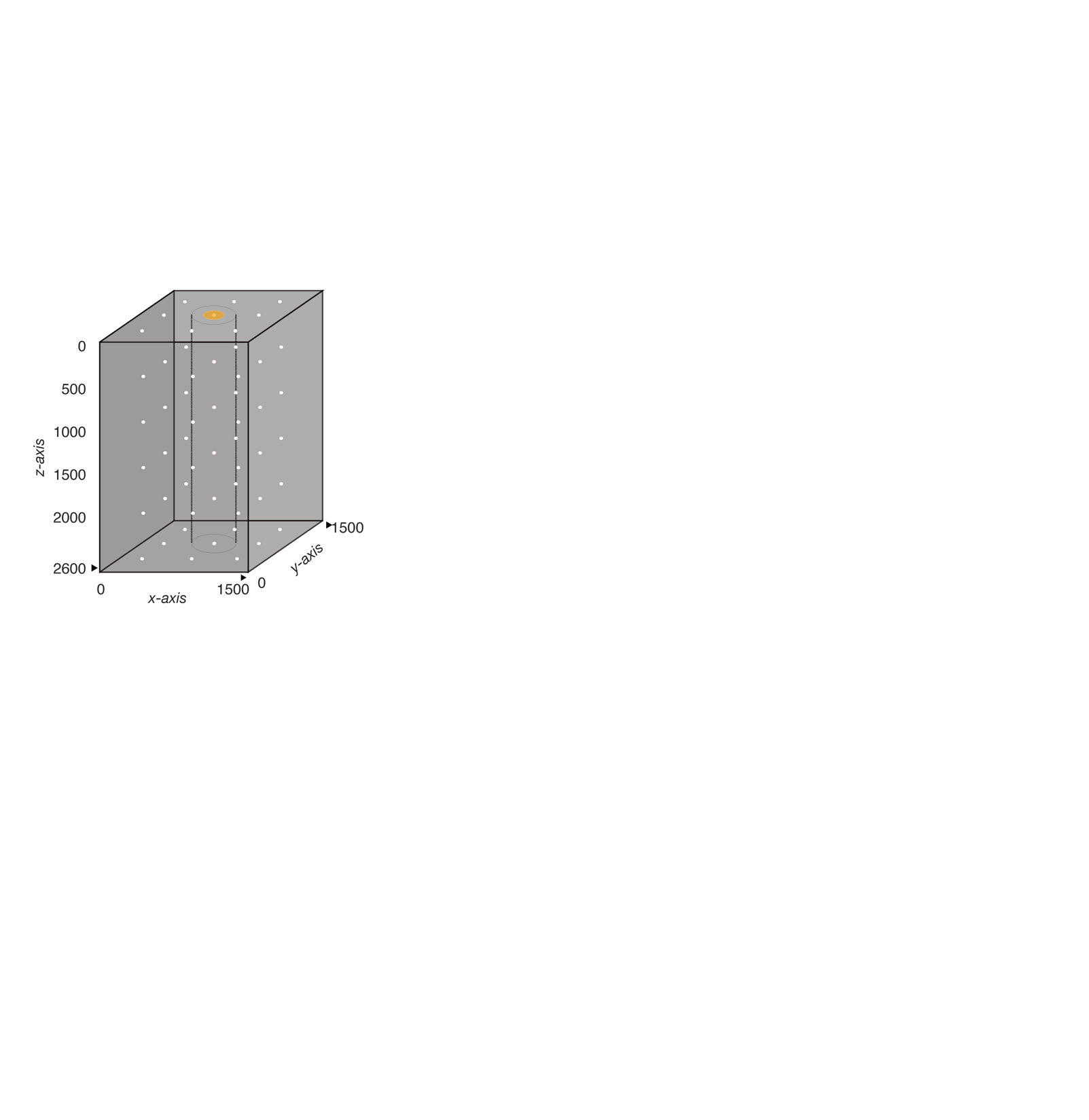

*Supp Fig. 9 Simulation architecture for neuron spiking model. Electrode recording sites are shown as white points. Stimulation site is shown as a yellow dot on the surface of the simulated tissue. The vertical cylindrical column defines the region directly below our simulated illumination. Neurons in this portion of the tissue have their raster plots shown in Supp. Fig. 5a.*

### Supplemental Tables

Supp. Table 1: Total successful behavioral trial counts.

|  | Monkey H Trials, Stimulation | Monkey H Trials,  No stim | Monkey L Trials, Stimulation | Monkey L Trials,  No stim |
| --- | --- | --- | --- | --- |
| Right | 554 | 570 | 1275 | 1356 |
| Down | 588 | 544 | 1389 | 1556 |
| Left | 579 | 547 | 1404 | 1450 |
| Up | 560 | 580 | 1391 | 1280 |
| Total | 2281 | 2241 | 5459 | 5642 |

Supp. Table 2: Statistics for behavioral metrics. Two-sample t-test was used to compare differences in medians of reach time and path length.

| P values | **Monkey H** | | **Monkey L** | |
| --- | --- | --- | --- | --- |
|  | Reach Time | Path Length | Reach Time | Path Length |
| **Right** | 0.13 | 0.36 | 0.98 | 0.82 |
| **Down** | 0.03 | 0.02 | 2.0 x 10^-4^ | 2.6 x 10^-10^ |
| **Left** | 0.22 | 0.35 | 0.03 | 1.4 x 10^-5^ |
| **Up** | 0.99 | 0.21 | 0.33 | 0.63 |
| **Combined** | 0.04 | 0.31 | 8.2 x 10^-3^ | 8.8 x 10^-4^ |

Supp. Table 3: Electrode count and trial numbers for power analysis shown in manuscript Fig. 6.

|  | **Monkey H** | | **Monkey L** | | | **Monkey C** |
| --- | --- | --- | --- | --- | --- | --- |
|  | Red | Red 2 years after injection | Red | Red with tissue growth | Blue | Red |
| **Opsin expressing electrode count** | 9 | 10 | 12 | 6 | 12 | 0 |
| **Opsin deficient electrode count** | 3 | 20 | 9 | 3 | 9 | 28 |
| **Trial #** | 30 | 60 | 100 | 50 | 100 | 100 |

### Supplemental Video

Supp. Vid. 1: Demonstration of LED illumination with spatiotemporal complexity and dynamic optical power. Note the final flash is from only 4 LEDs but is at high power and washes out the video camera.

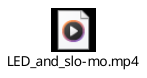
